# Sexual antagonism in sequential hermaphrodites

**DOI:** 10.1101/2023.03.01.530720

**Authors:** Thomas J. Hitchcock, Andy Gardner

## Abstract

Females and males may have distinct phenotypic optima, but share essentially the same complement of genes, potentially leading to trade-offs between attaining high fitness through female versus male reproductive success. Such sexual antagonism may be particularly acute in hermaphrodites, whereby both reproductive strategies are housed within a single individual. Whilst previous models have focused on simultaneous hermaphroditism, we lack theory for how sexual antagonism may play out under sequential hermaphroditism, which have the additional complexities of age-structure. Here we develop a formal theory of sexual antagonism in sequential hermaphrodites. First, we develop a general theoretical overview of the problem, then consider different types of sexually antagonistic and life-history trade-offs, under different modes of genetic inheritance (autosomal or cytoplasmic), and different forms of sequential hermaphroditism (protogynous, protoandrous or bi-directional). Finally, we provide a concrete illustration of these general patterns by developing a two-stage two-sex model, which yields conditions for both invasion of sexually antagonistic alleles and maintenance of sexually antagonistic polymorphisms.

## Introduction

Females and males may have distinct phenotypic optima, encompassing different behaviours, morphologies, and physiologies (Darwin 1871; Parker 2014). However, the sexes, for the most part, share an identical complement of genes, and this can lead to trade-offs between attaining high fitness through each of these two routes, with alleles that are beneficial in relation to female reproductive success being costly in relation to male reproductive success, and vice versa. Whilst the dynamics of sexual antagonism have long been studied (Owen 1953; Parsons 1961; Kidwell et al. 1977; Rice 1984), this was typically in a restricted set of conditions (e.g. panmixia, non-overlapping generations), and there has been an increased recent interest in how sexual antagonism may be expected to play out in a more diverse range of ecological and genetic scenarios. This has included: population structure and inbreeding (Jordan and Connallon 2014a; Tazzyman and Abbott 2015; Hitchcock and Gardner 2020; Flintham et al. 2021; Hitchcock et al. 2021), unusual modes of inheritance (Jaquiéry et al. 2013; Hitchcock et al. 2021; Klein et al. 2021), age structure (de Vries and Caswell 2019; Hitchcock and Gardner 2020), and mechanisms of population regulation (Harts et al. 2014; Connallon et al. 2019, de Vries and Olito 2022).

One group of organisms within which sexual antagonism may be particularly acute are hermaphrodites. In these organisms female and male reproductive strategies are housed within a single individual, and thus they may experience more intense sexual antagonism than do dioecious species (Abbott 2011). Previous theoretical work considering sexual antagonism in hermaphroditic organisms has primarily focused on simultaneous hermaphrodites—i.e. when individuals produce sperm and eggs concurrently throughout their life (Jordan and Connallon 2014b; Tazzyman and Abbott 2015; Olito 2017; Olito et al. 2018; de Vries and Olito 2022). In contrast, sequential hermaphroditism—i.e. when individuals temporally separate the production of male and female gametes—has been relatively overlooked. Although there have been some verbal treatments (Abbott 2011; Schärer et al. 2015; Abbott et al. 2019), no formal theory has yet been developed on sexual antagonism under sequential hermaphroditism.

Yet sequential hermaphrodites exhibit many interesting features that may make them exceptional study systems for improving our understanding of sexual antagonism. Firstly, due to the way in which female and male strategies are temporally stratified, age structure is likely to be an important modulator of evolutionary outcomes. Recent work has shown how sex differences in fecundity and mortality schedules may shape sexual antagonism in dioecious (gonochoristic) taxa (de Vries and Caswell 2019; Hitchcock and Gardner 2020), and thus sequential hermaphrodites, with their diversity of combinations of age and sex structure—from protandry (male first), to protogyny (female first), and bidirectional sex change (Leonard 2018)—may prove superb systems with which to test these principles.

Moreover, whilst in dioecious systems the flow of genes between sexes occurs solely through reproduction, in sequential hermaphrodites this also occurs through sex change. This may have profound consequences. For instance, males are typically an evolutionary dead end for mitochondrial genes in dioecious species, as they usually do not pass these genes onto their offspring, and mitochondria are therefore liable to accumulate male-deleterious mutations—the “mother’s curse” (Frank and Hurst 1996; Gemmell et al. 2004). In contrast, a male’s mitochondria will have reproductive value insofar as he is able to change sex and produce offspring as a female later in life, such that male-deleterious mitochondrial alleles need not be invisible to selection in sequential hermaphrodites.

Here we develop a formal theory of sexual antagonism in sequential hermaphrodites. We construct a general theoretical framework, which reveals how the reproductive values of males and females are shaped by transmission genetics, age structure, and patterns of sex change, and we use this to investigate how different sexually antagonistic and life-history trade-offs are modulated by differences in the mode of genetic inheritance (autosomal or cytoplasmic) and the form of sequential hermaphroditism (protogynous, protoandrous or bi-directional). We provide a concrete illustration of these general patterns by developing a two-stage two-sex model, yielding conditions for both the invasion of sexually antagonistic alleles and the maintenance of sexually antagonistic polymorphisms.

### Reproductive value in age and sex-structured species

When populations are divided into different classes, such as age and/or sex, then the relative force of selection (Hamilton 1966)—and thus weights upon selective effects in these different classes—are given by their reproductive values, i.e. the expected asymptotic fraction of genes in future generations that will descend from that particular class of individuals in the current generation (Fisher 1930; Price and Smith 1972; Taylor 1990, 1996; Grafen 2006). If transitions between classes are governed by an aperiodic Markov process that has attained its steady state behaviour (i.e. stable class distribution), then these contributions to the future are equivalent to the relative amounts of time that a gene’s lineage will have spent in a particular state. If we assume that all newborns have the same independent probability of being male or female, then we may calculate a class’s reproductive value simply by sampling a gene from a newborn, tracing its origin back to one of the newborn’s parents, and determining the total extent of the parent’s life up to this moment that was spent as a member of this class—and then taking the expectation of this quantity across all newborns (further derivation can be see in SM§2.4).

Sampling a gene from a newborn individual, we notate the probability that it derived from the mother by *ζ*, and the probability it derived from the father by 1 – *ζ*. If it derived from the mother, then with probability *μ_a_* she is of age *a*, and if derived from the father, then with probability *v_a_* he is of age *a*, where ∑*_a_ μ_a_* = ∑*_a_ v_a_* = 1 and the units of time are entirely arbitrary. For an age-*a* mother of a newborn, let *x_a_* be the proportion of her life that, on average, she has spent as a female and, for an age-*a* father or a newborn, let *y_a_* be the proportion of his life that, on average, he has spent as a male. The long-term proportion of time spent in males and females—and thus the ratio of reproductive values (*c*_f_/*c*_m_)—will then be given by:

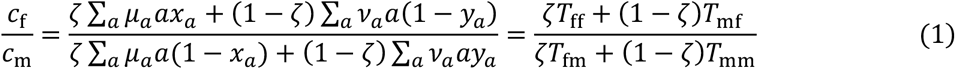

where *T*_ij_ is the mean time that the sex-i parent of a newborn has spent as sex j.

For example, in a simple dioecious species, *x_a_* and *y_a_* are simply 1, i.e. individuals spend their whole lives as the sex that they were born, and therefore *T*_fm_ = *T*_mf_ = 0. This means that in a dioecious species the ratio of average time that a gene has spent as a female versus average time spent as a male through the generations is given by:

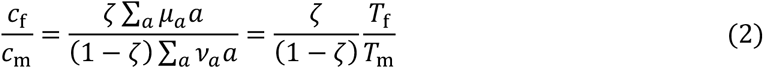

where *T*_f_ is the mean maternal age, and *T*_m_ is the mean paternal age (Grafen 2015, Hitchcock and Gardner 2020).

Inspecting these equations, we can gauge some general patterns relating different forms of sequential hermaphroditism to the reproductive values of females and males. For instance, if species are protogynous, then this will mean that fathers will have spent some time as females (*T*_mf_ > 0), whilst mothers will not have spent any time as males (*T*_fm_ = 0), which—all else being equal—serves to increase the reproductive value of females. In contrast, under protandry, mothers will have spent a fraction of their life as males (*T*_fm_ > 0), whilst fathers will not have spent any time as females (*T*_mf_ = 0), which—all else being equal—serves to increase the reproductive of males.

We can further partition the reproductive values of these classes into the reproductive values of—and thus force of selection acting upon—different types of class transition. To compute these, we first need to describe the frequency of each type of class transition. For instance, the frequency of the transition from a parent of a given sex to the newborn class (i.e. the value of reproduction for that sex), will be given by the frequency of the newborn class, and the probability that genes in a newborn trace back to the parent of the given sex. As a gene passes through the generations, this newborn class, will, on average, be encountered every 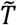 time units, where 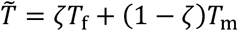. This quantity, equivalent to a weighted mean parental age, is arguably the most natural measure of generation time, and when defined as thus gives the result that the reproductive value of newborns is simply 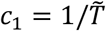 (recovering the result for asexuality of Bienvenu and Legendre (2015)). As this class comes from females with probability ζ, and from males with probability 1 – *ζ*, then the relative reproductive value of reproduction through the two sexes is given by:

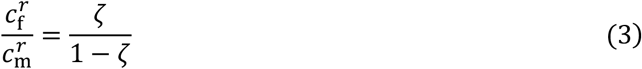

This mirrors previous results which show that it is the transmission genetics alone – and not the demography – which shape the relative ancestral contribution through reproduction made by males and females (Grafen 2014; Hitchcock and Gardner 2020). We may similarly calculate the reproductive value of survival by males and females, the ratio of which is given by:

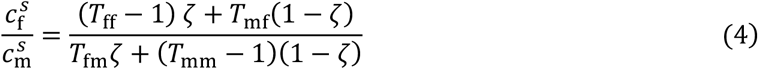

Once again, if we assume that there is no sex change (*T*_mf_ = *T*_fm_ = 0), then we recover the results of Hitchcock and Gardner (2020) as a special case 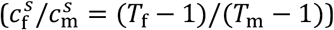. Having obtained these reproductive values, we can now investigate the consequences of trade-offs between these different states and state-transitions, and, in particular, trade-offs between males and females, i.e. sexual antagonism. We first investigate the consequences for autosomal genes (*ζ* = 1/2), before considering these same trade-offs as experienced by cytoplasmic genes (*ζ* ≈ 1).

### Survival and fecundity trade-offs

First, we consider a sexually antagonistic allele that solely affects fecundity. This allele confers a marginal fecundity benefit of *σ* upon one sex, and a marginal fecundity cost of *τ* upon the other (Table 1). Assuming weak selection, no population structure, and no assortative mating, then the condition for a female-beneficial allele to invade is 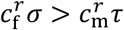, and for a male-beneficial allele it is 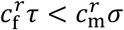. We may then rewrite the condition for a female-beneficial allele to invade as *τ/σ* < *F*, and the condition for a male-beneficial allele to invade in the form *τ/σ* < 1/*F*, where *F* describes the ‘potential for feminisation’ (cf. Hitchcock et al. 2021). If *F* > 1, then the condition for invasion is less stringent for female-beneficial alleles than for male-beneficial alleles. Conversely if *F* < 1, then the condition for invasion is more stringent for female-beneficial alleles than for male-beneficial alleles. Some rearrangement reveals that for autosomal genes:

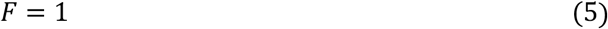

Such that under arbitrary age structure and patterns of sex change, the invasion condition is equally stringent for male-beneficial and female-beneficial alleles.

**Table 1:** The relationship between the fitness schemes used to plot the invasion conditions in Figure 1, taken from Kidwell (1977), and the marginal fitness effects (*σ, −τ*) discussed in the main text. Marginal fitness effects are calculated in the limit of weak selection, and when the allele is vanishingly rare in the population.

|  | Females | Males |
| --- | --- | --- |
| Genotypic fitnesses |  |  |
| $W_{00}$ | 1 | $1 - s_m$ |
| $W_{01}/W_{10}$ | $1 - h_f s_f$ | $1 - h_m s_m$ |
| $W_{11}$ | $1 - s_f$ | 1 |
| Marginal fitness effects |  |  |
| $\sigma$ | $(1 - h_f)s_f$ | $(1 - h_m)s_m$ |
| $\tau$ | $h_f s_f$ | $h_m s_m$ |

Next, we consider a sexually antagonistic allele that solely affects survival. Similar to above, this allele confers a marginal survival benefit of *σ* upon one sex, and a marginal survival cost of *τ* upon the other. Under the same assumptions as above, the condition for invasion will be 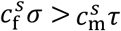 for a female-beneficial allele, and 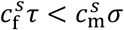 for a male beneficial allele. We can once again rearrange the invasion conditions into a potential for feminisation where 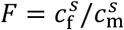. Substituting in the appropriate reproductive values from above, the potential for feminisation becomes:

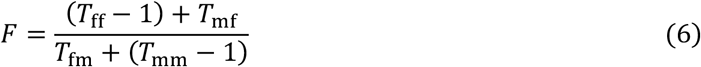

As well as intersexual trade-offs between fitness components, we may envisage trade-offs between different fitness components between males and females. First, we consider a trade-off between female fecundity and male survival. In this case, the condition for a female-beneficial allele to invade is 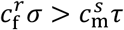, and for a male beneficial allele it is 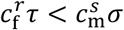. Again, substituting in the reproductive values, the potential for feminisation becomes:

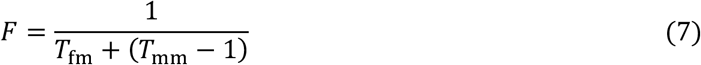

We may also consider an allele which modulates a trade-off between female survival and male fecundity. In this case the conditions for a female-beneficial and male beneficial allele will be 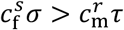 and 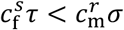 respectively. With the resultant potential for feminisation being:

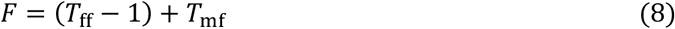

Alongside trade-offs between sexes, we may consider trade-offs that occur within a sex. If an allele confers a fecundity benefit *σ* but a survival cost *τ*, then if it is active in females it will invade provided 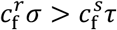 and if it is active in males it will invade provided 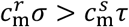.

For the purpose of concreteness and illustration, we additionally analyse a simple population genetic model of sexual antagonism that incorporates two possible age-classes (*a* ∈ {1,2}) and two possible sex-classes (*k* ∈ {f, m})—see Figure S1 for a visual representation of the life cycle and the associated life cycle parameters. Within this structure, we allow for a variety of sex change systems by describing the following parameters: *α* is the fraction of age-2 who females came from age-1 females in the previous generation, *β* is the fraction of age-2 males who came from age-1 males in the previous generation, *μ* is the fraction of newborns who have age-1 mothers, and *ν* is the fraction of newborns who have age-1 fathers. Thus, simple dioecy can be recovered by setting *α* = *β* = 1, simple protandry can be recovered by setting *α* = 0 and *β* = 1, and simple protogyny can be recovered by setting *α* = 1 and *β* = 0. Using this model we may compute the various invasion conditions for different types of sexually antagonistic trade-offs in terms of the above parameters, results of which are plotted in Figure 1, with the associated fitness scheme given in Table 1. Additionally, expressions for the potential for feminisation for various combinations of these trade-offs are given in Table 2.

**Figure 1:**
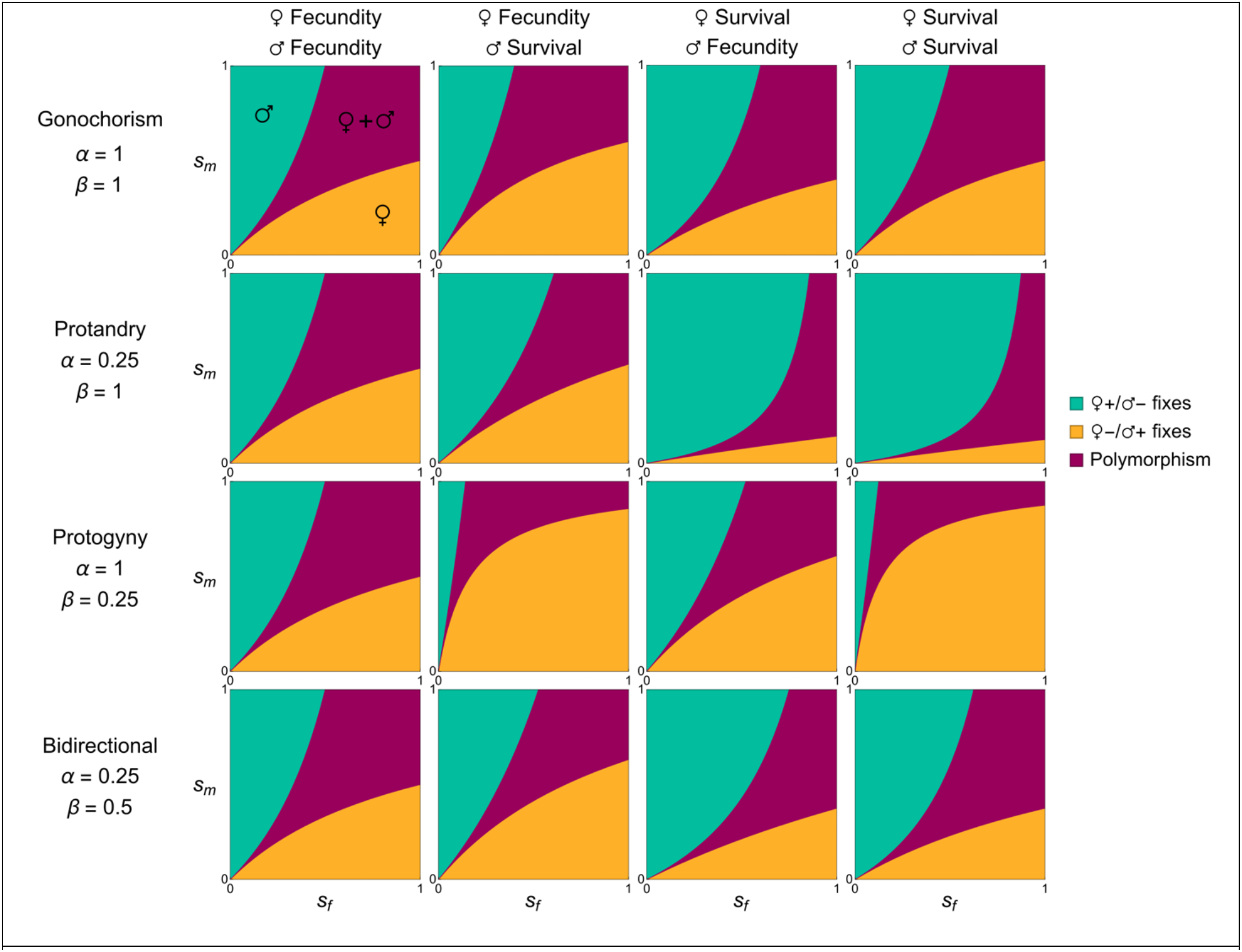
Invasion conditions for an allele affecting various types of sexually antagonistic trade-off under equal dominance. Trade-offs plotted include: fecundity in both sexes, fecundity in females and survival in males, survival in females and fecundity in males, and survival in both sexes. And for various types of sexual system including dioecy, protandry, protogyny and bidirectional sex change. The meaning of key parameters are described in Table 2 and Figure S1. For all plots we assume that *μ* = *v* = 1/3, and *h_f_* = 1 – *h_m_*.

**Table 2:** The potential for feminisation *F*, under different types of sexually antagonistic trade-offs in the two-stage model, whose life cycle is illustrated in Figure S1. *ζ* is the probability a gene copy in a juvenile was inherited from a female, *α* is the probability a gene copy in an age-2 female came from an age-1 female. *β* is the probability a gene copy in an age-2 male came from an age-1 male. *μ* is the fraction of mothers from the age-1 class, *v* is the fraction of fathers from the age-1 class.

|  | Male fecundity | Male survival |
| --- | --- | --- |
| Female fecundity | $\frac{\zeta}{(1 - \zeta)}$ | $\frac{\zeta}{\zeta(1 - \mu)(1 - \alpha) + (1 - \zeta)\beta(1 - \nu)}$ |
| Female survival | $\frac{\zeta\alpha(1 - \mu) + (1 - \zeta)(1 - \nu)(1 - \beta)}{\zeta(1 - \zeta)}$ | $\frac{\zeta\alpha(1 - \mu) + (1 - \zeta)(1 - \nu)(1 - \beta)}{\zeta(1 - \mu)(1 - \alpha) + (1 - \zeta)\beta(1 - \nu)}$ |

### Cytoplasmic genes

Thus far we have considered only autosomal genes. However, a range of important functions are encoded in genomes outwith the nucleus, for example in chloroplasts, mitochondria, and other symbionts (Birky 2001; Camus et al. 2022). Moreover, such elements are interesting from the perspective of sexual antagonism as—due to their often strictly matrilineal inheritance, and a concomitant absence of selection in relation to their male carriers—they are liable to accumulate mutations that are deleterious to males (Frank and Hurst 1996; Gemmell et al. 2004), with the converse argument applying to those cytoplasmic elements which show a primarily patrilineal mode of inheritance. The logic of this “mother’s curse” (and “father’s curse”) has not been explored in relation to sequential hermaphrodites, wherein such cytoplasmic genes currently residing in males may still contribute to future generations even if they are never transmitted via male reproduction. Here, we investigate a broad range of cytoplasmic transmission modes, by considering an element which is inherited with probability 1 – *ζ* from a father.

Returning to our earlier invasion condition 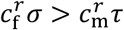 for a (female-beneficial) sexually antagonistic allele affecting fecundity, the potential for feminisation is:

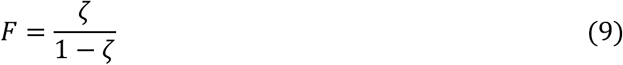

Assuming that individuals have only one cytoplasmic type (Kuijper et al. 2015), then, in the limit of full matrilineal inheritance (i.e. *ζ* = 1), the potential for feminization for a cytoplasmic gene is *F* = +∞. Thus, as with conventional dioecious organisms (and other hermaphrodites), if there is strict matrilineal inheritance then cytoplasmic genes place no value upon male fecundity relative to female fecundity.

In contrast, the potential for feminisation for a sexually antagonistic allele affecting survival is:

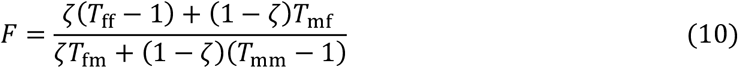

which, in the limit of strict matrilineal inheritance (*ζ* = 1), becomes *F* = (*T*_ff_ – 1)/*T*_fm_. Thus, the in such a case the relative force of selection upon survival in males, relative to females, for a cytoplasmic gene ultimately becomes the average time spent in that state during a reproductive female’s lifetime. This arises because even with strict matrilineal inheritance, provided there is sex change from male to female, then male survival prior to sex change may ultimately contribute to female reproduction, and thus future generations. This leads to some striking asymmetries between different types of sequential hermaphroditism. For protogynous species, whereby individuals are female first and male later (*T*_fm_ = 0), cytoplasmic genes will place no value upon male survival *F* = +∞. In contrast, for protandrous species, whereby individuals are male first and female later (*T*_fm_ > 0), cytoplasmic genes may place value upon male survival, even at a cost to the survival of females.

For trade-offs between female fecundity and male survival the potential for feminisation is:

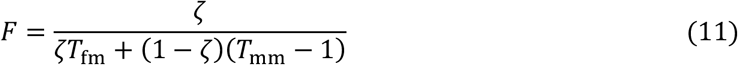

Thus, as before, for a protogynous species and strict matrilineal inheritance, cytoplasmic genes place no value upon males, such that *F* = +∞, whilst for a protandrous species male survival is of value, such that *F* = 1 /*T*_fm_. And for trade-offs between female fecundity and male survival the potential for feminization is:

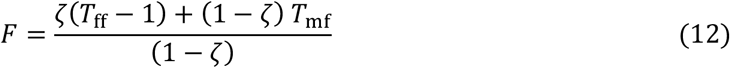

Thus, once again for a matrilineally inherited cytoplasmic gene (*ζ* = 1) the potential for feminisation is *F* = +∞, as no genes are passed on via male reproduction. We also consider some of the within sex life-history trade-offs in SM§3.5.

## Discussion

Here we have outlined how the reproductive values of—and thus the force of selection upon—females and males are altered by different systems of sequential hermaphroditism, and thus how different sexually antagonistic trade-offs may manifest differently in these organisms as compared to dioecious species. We have shown that, regardless of the system of sex change or other sex-specific demographic parameters, the force of selection upon fecundity effects of males and females is equal—for autosomal genes—due to their equivalent contributions to the newborn class. However, trade-offs involving survival may not be, either because the mean parental age of one sex is higher than the other, but also because, with the possibility of sex change, one sex may contribute (more) to the other through survival. This results in distinct patterns of sexually antagonistic trade-offs involving survival with different forms of sequential hermaphrodites (e.g. protandry versus protogyny), and in ways that are distinct from dioecious species. Moreover, this flow of genes that occurs between the sexes as a consequence of sex-change means that cytoplasmic genes, which are normally strictly matrilineally inherited, may nevertheless experience selection in males, and this yields clear-cut predictions concerning comparisons between dioecious species and different forms of sequential hermaphrodism.

Whilst recent years have seen a renaissance in the development of theory on sexual antagonism, there has—until now—been a lack of any formal theory regarding sexual antagonism in relation to sequential hermaphrodites (Abbott 2011; Schärer et al. 2015). And, although it has been suggested that trade-offs might be biased towards the first sex due to a skewed sex ratio (Abbott et al. 2019), this issue has remained unresolved. Here we have shown more generally how patterns of sex-change—varying from protandry to protogyny to bidirectional sex change—may generate biases in the force of selection towards one sex or the other, and how this varies across different fitness components (e.g. survival versus fecundity). As these patterns depend upon the direction of sex change and other demographic parameters, sequential hermaphrodites provide excellent comparative testbeds for the theory of sexual antagonism more generally, just as they have for the theory of sex allocation (Charnov 1982; West 2009). Indeed, many systems which have previously been investigated in relation to sex allocation may also prove good systems to understand sexual antagonism. For instance, the Pandalid shrimp (*Pandalus jordani)* has long been studied with regards to sex allocation (Charnov 1979; Charnov and Hannah 2002; Charnov and Groth 2019). This protandric species typically has just two breeding seasons, and shows variation amongst populations in the number of males who change sex, in part responding to differential harvesting pressure that alters the age distribution of the local population. These subpopulations may well show variation in both mean maternal and paternal ages, as well as variation in the timing and extent of sex change, and thus provide good systems for testing our theoretical predictions.

Not only is there little current theory regarding sexual antagonism in sequential hermaphrodites, but currently also a relative lack of empirical work (Abbott 2011; Schärer et al. 2015). This may stem from there being very few ‘classical’ molecular model organisms that are sequential hermaphrodites, although new genomic tools have made molecular investigation of such non-models organisms more tractable. Combined with new theoretical developments (Cheng and Kirkpatrick 2016; Connallon and Matthews 2019; Ruzicka et al. 2020; Ruzicka and Connallon 2022; Ruzicka et al. 2022; Lucotte et al. 2022), detection of sexually antagonistic alleles—and the signatures of sexually antagonistic selection—may be increasingly feasible in these groups. For instance, the genome of the protandrous gilthead sea bream (*Sparus aurata)* has recently been sequenced, and its sex-biased gene expressions described (Pauletto et al. 2018), and this may make this species, as well as other Sparids, good candidates for future investigation. Additionally, one example that has previously been suggested as representing sexual antagonism is aggression in the protogynous sharpnose sandperch (*Parapercis cylindrica)* (Sprenger et al. 2012; Schärer et al. 2015). Here, individuals who are aggressive as females also tend to be aggressive as males later in life. However, the sex-specific optima in relation to aggression was assumed rather than demonstrated, so further investigation of other traits—such as colouration, physiology, and behaviour—which are known to have sex-specific optima may be worthwhile to see whether these too reveal such correlations.

Alongside sexual antagonism, the theory we have outlined here also provides a framework with which to understand the forces shaping other life-history trade-offs, in particular senescence. The interplay between sex and senescence has recently received increasing theoretical and empirical attention (Maklakov and Lummaa 2013; Adler and Bonduriansky 2014; Marais and Lemaître 2022; Bronikowski et al. 2022) and, once again, sequential hermaphrodites provide a particularly interesting set of species and life cycles with which to investigate these phenomena, as questions about the relative value of different age classes are intrinsically bound up with questions about the value of different sexes. One particular suggestion has been that protogynous species with extreme breeding sex ratios may experience reduced senescence as, in such species, a relatively old individual might still have a great deal of reproductive potential (Charnov 1993). More generally, it is interesting to consider whether, if half of the future reproduction is forced to come from above a certain age, this will shift the force of selection to relatively later in life. Conversely, it is unclear whether the reverse argument may also apply, and that by constraining much (or all) of one sex’s reproduction to younger ages, then the force of selection upon older age classes is reduced relative to a comparable dioecious species. Future modelling is needed to address these questions and ensure that empirical work is best targeted to the most salient comparisons.

We have also shown how sequential hermaphroditism will alter patterns of selection in relation to maternally-inherited cytoplasmic genes. Whilst it has long been understood from a theoretical perspective that cytoplasmic and autosomal genes may differ in their interests when there are trade-offs between the sexes, e.g. “mother’s curse” (Frank 1996, Gemmell 2004), more recently there has been an increased amount of empirical support for this effect (Nagarajan-Radha et al. 2020; Carnegie et al. 2021). We have shown here that—unlike in dioecious species—cytoplasmic genes may undergo direct selection in males, with the extent of this depending on the form of hermaphroditism considered, leading to potentially neat empirical tests. Moreover, cytoplasmic genes are expected to make different trade-offs regarding the pace of life as compared to autosomal genes. For instance, in protogynous species, mitochondria do not reproduce through the second sex, and should thus favour a faster pace of life than autosomal genes. In contrast, in protandrous species, mitochondria are expected to place a relatively greater value on reproduction later in life, and thus favour slower life histories than autosomal genes. Such intragenomic conflicts of interest are expected to be strong, and these are phenotypes for which mitochondria are likely to play important modulating roles (Hill 2019).

Moreover, such cytoplasmic genes would be expected to favour not only a different pace of life to autosomal genes, but also a distinct allocation to male and female strategies. This has previously been well studied in the form of cytoplasmic male sterility in hermaphrodites (especially plants)(Frank 1989; Chase 2007; Kim and Zhang 2018) and male-killing endosymbionts in dioecious species (especially arthropods)(Hurst 1991; Massey and Newton 2022; Hornett et al. 2022). Yet, despite the centrality of sequential hermaphrodites to sex-allocation research, little empirical or theoretical work has investigated the potential for such sex allocation distorting elements in sex-changing species. These species may potentially reveal distinct mechanisms of sex-ratio distortion, potentially either by delaying or preventing sex change in protogynous species or favouring either earlier sex change or altering the sex at birth in protandrous species.

Here we considered conditions for invasion when fitness effects are vanishingly small. Typically, weak-selection assumptions do not matter greatly for sexual antagonism on autosomal genes (at least with non-overlapping generations), the reason being that—even under strong selection—the distribution of a sexually antagonistic allele across classes is not affected by selection in the previous generation, as alleles are reshuffled between the sexes every generation (hence issues detecting such alleles (Kasimatis et al. 2019)). However, this will not happen for sequential hermaphrodites because—just as in age-structured populations in asymmetric genetic systems—strong selection is liable to perturb the distribution of the mutant allele across classes (Abbott 2011; Schärer et al. 2015). This second-order effect may potentially alter the invasion conditions for sexually antagonistic alleles, making them either more—or less—stringent depending on whether strong selection generates negative or positive associations with the class to which it confers benefits, altering the potential for polymorphism in these groups. Moreover, these patterns may well differ amongst different types of sequential hermaphrodites. For instance, a sexually antagonistic allele which confers fecundity benefits to females in a protogynous species will become relatively more abundant in the newborn (female) class of individuals, amplifying its benefits, whilst conversely a male-beneficial allele will also become more abundant in newborn females, thus dampening its benefits. Future modelling should address this to both understand these effects in and of themselves, and to identify the systems and scenarios wherein weak selection approximations may be less accurate.

We have also focused on a relatively abstract model of sex change, in which individuals are indexed solely by age and sex. This, however, ignores many of the complexities that are associated with the ecologies that shape sex change and social interactions in these groups. For instance, in many species sex change is a highly social affair, with its timing governed by the local sex ratio, and an individual’s relative age, size, and condition (Charnov 1982; West 2009; Leonard 2018, Benvenuto and Lorenzi 2023). In the polychaete worm *Ophryotrocha puerilis* for instance, individuals are socially monogamous, and throughout the breeding season a pair of individuals may change sex repeatedly, with the larger individual being the female and the smaller the male, changing sex as their relative size changes (Berglund 1986). This also may provide a nice species for comparisons as closely related species are both hermaphroditic and dioecious (Dahlgren 2001). Moreover, alongside social interactions determining sex change, such sexually antagonistic fitness effects may also manifest through social interactions, and thus have more complex dynamics if they involve either siblings (e.g. sib-cannibalism in *Crepidula coquimbensis* (Brante et al. 2016)), or mates (e.g. sexual conflict (Schärer et al. 2015)). Additionally, particular systems may involve more complicated class structure. For example, in the protandrous slipper limpet (*Crepidula fornicata)* individuals can store sperm, such that females may produce offspring with previous mating partners who exist as females at the time of the offsprings’ conception (Broquet et al. 2015). In these cases, sperm themselves must be separately tracked as a separate class in order not to incorrectly assign those fitness effects to males who may either have died or else become females. Building in such details will help tailor future models to particular taxonomic groups, as well as developing a better understanding of which ecological factors may drive differences in the time that genes spend in male and female bodies, the magnitude of fitness effects they may impose and different points in the lifecycle, and thus which factors and systems may be best to focus on for experimental or comparative studies of sexual antagonism.

Finally, the models we have outlined here have focused on cases whereby, although an individual may reproduce as either sex throughout their life, at any given point in time they are exclusively reproducing as either a female or a male. However, in reality this boundary between sequential and simultaneous hermaphroditism is more porous (Leonard 2018). For instance, in both Lysmata shrimp (Baeza 2008) and certain polychaete worms (Sella 1990) individuals are protandric simultaneous hermaphrodites, reproducing as males when young, and then hermaphrodites when older. Additionally, many simultaneous hermaphrodites alter their sex allocation strategy as a function of their age, size, condition, and environmental variables and thus these too can be viewed as existing somewhere on the spectrum between strict sequential and strict simultaneous hermaphroditism (Leonard 2018). These examples further blur the boundary between questions of sex allocation, sexual antagonism, and life-history trade-offs, and thus require richer theory to help us identify and understand the ecological factors moulding the shape and pattern of life across these diverse sexual systems.

## Acknowledgements

We thank S. Immler, R. Iritani, G. Ruxton, V. Litzke, and K. Stucky for helpful discussion.

