## Supplementary Material for "Sexual antagonism in sequential hermaphrodites"

#### 1 Summary

The supplementary material is composed of two sections. In Part I, the distribution of reproductive values through the life-cycles is described. We do this for an asexual, hermaphrodite (simultaneous and sequential), and gonochorist case. In Part II, a concrete population genetics model of sexual antagonism is considered, where the conditions for invasion and polymorphism of both sexually antagonistic, and antagonistically pleiotropic alleles are considered.

#### 2 Life cycles and class reproductive values

##### 2.1 Reproductive values and the force of selection

In this section, we describe how reproductive value is distributed through various types of life-cycle. Reproductive value describes the expected fraction of the ancestry of an asymptotic population that descend from a gene copy (or set of gene copies, whether that be an individual or a class) in the current population (Taylor, 1990; Grafen, 2006). Moreover, these reproductive values also correspond to the force of selection upon the relative marginal fitness effects in these different classes (Taylor and Frank, 1996; Frank, 1998), and are equivalent to the elasticities with respect to population growth (Bienvenu and Legendre, 2015; Giaimo, 2022).

In the cases we analyse, we assume that the population has attained its stable age distribution, and that the various parameters of interest (and subsequent reproductive values) do not change as a function of time, and that the various probabilities of jumping between states depend only on the state of interest (i.e. is a Markovian process). To calculate the class reproductive values $c(i)$, we describe the trajectory of a gene lineage backwards in time. We consider the mean time of first return of class $i$, $\mathscr{R}(i)$.The inverse of this quantity is the stationary distribution ($c(i) = 1/\mathscr{R}(i)$), which simultaneously describes the frequency with which a gene state is visited in the ancestry of the population, as well as the expected contribution of that gene state to future populations, i.e. the class reproductive value.

#### 2.2 Asexual reproduction

First, we consider the asexual case. Let us define a function $\phi(x)$, which describes the probability that a gene lineage currently in the newborn state jumps to an individual of age $x$ in the next time step. We will focus on the case of discrete time, although this may readily be extended to continuous time. For a newborn individual, the mean return time will be:

$$\mathscr{R}(1) = \sum_{x=1}^{\infty} \phi(x)x = T \tag{S1}$$

This is also equivalent to the mean parental age or generation time $T$ (Bienvenu and Legendre, 2015). We now calculate the mean return time for other ages. To do this, let's define the probability that a gene lineage 'jumps' from the newborn state to age $a$ or higher:

$$\gamma(a) = \sum_{x=a}^{\infty} \phi(x) \tag{S2}$$

Let's define the expected time of a "successful" loop through the life cycle (i.e. one that includes age $a$) to be $t_{\bullet}(a)$, and the expected time of an unsuccessful loop (i.e. one that doesn't include age $a$) to be $t_{\circ}(a)$. Where:

$$t_{\bullet}(a) = \frac{\sum_{x=a}^{\infty} \phi(x)x}{\sum_{x=a}^{\infty} \phi(x)} = \frac{T_{\bullet}(a)}{\gamma(a)} \tag{S3}$$

And:

$$t_{\circ}(a) = \frac{\sum_{x=1}^{a-1} \phi(x)x}{\sum_{x=1}^{a-1} \phi(x)} = \frac{T_{\circ}(a)}{1-\gamma(a)} \tag{S4}$$

The mean return time for age $a$ can be broken down into three portions. The time back to the newborn state, the expected number of failed jumps (i.e. jumps below $a$), and the successful jump.

$$\mathscr{R}(a) = a + \left(\frac{1-\gamma(a)}{\gamma(a)}\right) t_{\circ}(a) + (t_{\bullet}(a) - a) = t_{\bullet}(a) + \left(\frac{1-\gamma(a)}{\gamma(a)}\right) t_{\circ}(a) \tag{S5}$$

Simplifying this we get:

$$\mathscr{R}(a) = \frac{T_{\bullet}(a)}{\gamma(a)} + \left(\frac{1-\gamma(a)}{\gamma(a)}\right) \frac{T_{\circ}(a)}{1-\gamma(a)} = \frac{T}{\gamma(a)} \tag{S6}$$

And so:

$$c(a) = \frac{1}{\mathscr{R}(a)} = \frac{\gamma(a)}{T} \tag{S7}$$

As $\gamma(a)$ can be seen as the inverse cumulative distribution function of $\phi(x)$, then it is straightforward to show that:

$$\frac{\mathrm{d}c(a)}{\mathrm{d}a} \leq 0 \tag{S8}$$

i.e. the class reproductive value (and thus force of selection) never increases with age (Hamilton, 1966; Giaimo, 2022). In addition, we can partition the reproductive value into that through survival and that through reproduction. The probability of having entered the age $a$ through reproduction is:

$$c_r(a) = \phi(a)c(1) = \phi(a)/T \tag{S9}$$

And the probability of entering through survival is:

$$c_s(a) = c(a+1) = \gamma(a+1)/T \tag{S10}$$

#### 2.3 Sexual reproduction: simultaneous hermaphroditism

We can extend the logic of the above section to consider sexual reproduction. We first consider a simultaneous hermaphrodite. We now add the following notation: $\zeta$ is the probability that a gene copy in a newborn descends through an egg (i.e. through female reproduction), $1-\zeta$ the probability that it descends through sperm (i.e. male reproduction), $\phi_f(x)$ is the probability that - given a gene copy descends from a female - it comes from an individual of age $x$, and $\phi_m(x)$ is the probability that - given a gene copy comes from a male - it comes from a male of age $x$. Thus now:

$$T_f = \sum_{x=1}^{\infty} \phi_f(x)x \tag{S11}$$

$$T_m = \sum_{x=1}^{\infty} \phi_m(x)x \tag{S12}$$

Where $T_f$ and $T_m$ are sex-specific generation times. The mean return time for a newborn is thus:

$$\mathscr{R}(1) = \zeta T_f + (1-\zeta) T_m = \tilde{T} \tag{S13}$$

Once again, we can now consider other age classes. The probability of a "successful" loop for age $a$ will be:

$$\gamma(a) = \zeta \sum_{x=a}^{\infty} \phi_f(x) + (1-\zeta) \sum_{x=a}^{\infty} \phi_f(x) = \zeta\gamma_f(a) + (1-\zeta)\gamma_m(a) \tag{S14}$$

And the expected time of a successful loop will be:

$$\begin{aligned} t_\bullet(a) &= \left(\frac{\zeta\gamma_f(a)}{\gamma(a)}\right)\left(\frac{\sum_{x=a}^{\infty}\phi_f(x)x}{\sum_{x=a}^{\infty}\phi_f(x)}\right) + \left(\frac{(1-\zeta)\gamma_m(a)}{\gamma(a)}\right)\left(\frac{\sum_{x=a}^{\infty}\phi_m(x)x}{\sum_{x=a}^{\infty}\phi_m(x)}\right) \\ &= \left(\frac{\zeta\gamma_f(a)}{\gamma(a)}\right) t_\bullet^f(a) + \left(\frac{(1-\zeta)\gamma_m(a)}{\gamma(a)}\right) t_\bullet^m(a) \\ &= \left(\frac{\zeta\gamma_f(a)}{\gamma(a)}\right)\left(\frac{T_\bullet^f(a)}{\gamma_f(a)}\right) + \left(\frac{(1-\zeta)\gamma_m(a)}{\gamma(a)}\right)\left(\frac{T_\bullet^m(a)}{\gamma_m(a)}\right) \end{aligned} \tag{S15}$$

The expected time of an unsuccessful loop will be:

$$\begin{aligned} t_\circ(a) &= \left(\frac{\zeta(1-\gamma_{f)}(a)}{1-\gamma(a)}\right)\left(\frac{\sum_{x=1}^{a-1}\phi_f(x)x}{\sum_{x=1}^{a-1}\phi_f(x)}\right) + \left(\frac{(1-\zeta)(1-\gamma_m(a))}{1-\gamma(a)}\right)\left(\frac{\sum_{x=1}^{a-1}\phi_m(x)x}{\sum_{x=1}^{a-1}\phi_m(x)}\right) \\ &= \left(\frac{\zeta(1-\gamma_f(a))}{1-\gamma(a)}\right) t_\circ^f(a) + \left(\frac{(1-\zeta)(1-\gamma_m(a))}{1-\gamma(a)}\right) t_\circ^m(a) \\ &= \left(\frac{\zeta(1-\gamma_f(a))}{(1-\gamma(a))}\right)\left(\frac{T_\circ^f(a)}{1-\gamma_f(a)}\right) + \left(\frac{(1-\zeta)(1-\gamma_m(a))}{1-\gamma(a)}\right)\left(\frac{T_\circ^m(a)}{1-\gamma_m(a)}\right) \end{aligned} \tag{S16}$$

And so as before:

$$\begin{aligned} \mathscr{R}(a) &= t_\bullet(a) + \left(\frac{1-\gamma(a)}{\gamma(a)}\right) t_\circ(a) \\ &= \left(\frac{\zeta T_\bullet^f(a) + (1-\zeta) T_\bullet^m(a)}{\gamma(a)}\right) + \left(\frac{1-\gamma(a)}{\gamma(a)}\right)\left(\frac{\zeta T_\circ^f(a) + (1-\zeta) T_\circ^m(a)}{1-\gamma(a)}\right) \\ &= \tilde{T}/\gamma(a) \end{aligned} \tag{S17}$$

And so the class reproductive value of an age $a$ hermaphrodite.

$$c(a) = \gamma(a)/\tilde{T} \tag{S18}$$

Which we can once more partition into reproduction:

$$c_{r_f}(a) = \zeta\phi_f(a)c(1) = \frac{\zeta\phi_f(a)}{\tilde{T}} \tag{S19}$$

$$c_{r_\mathrm{m}}(a) = (1-\zeta)\phi_\mathrm{m}(a)c(1) = \frac{(1-\zeta)\phi_\mathrm{m}(a)}{\tilde{T}} \tag{S20}$$

And survival:

$$c_s(a) = c(a+1) = \gamma(a+1)/\tilde{T} \tag{S21}$$

#### 2.4 Sexual reproduction: sequential hermaphrodites

Now, we consider sequential hermaphrodites. Here, at each time point, individuals are either males or females, however they may change between these two states as they age. In addition to the above notation, we now introduction the following functions: $\mu_\mathrm{f}(x,y)$ is the probability that a gene lineage currently in an age $x$ female was in a female at age $y$, and similarly $\mu_\mathrm{m}(x,y)$ is the probability that a gene lineage currently in an age $x$ male was in an age $y$ male, in both cases $y \leq x$.

The probability of a successful loop for a female of age $a$ can be written as:

$$\gamma(f_\mathrm{a}) = \zeta\left(\sum_{x=a}^{\infty}\phi_\mathrm{f}(x)\mu_\mathrm{f}(x,a)\right) + (1-\zeta)\left(\sum_{x=a}^{\infty}\phi_\mathrm{f}(x)(1-\mu_\mathrm{m}(x,a))\right) = \zeta\gamma_\mathrm{f}(f_\mathrm{a}) + (1-\zeta)\gamma_\mathrm{m}(f_\mathrm{a}) \tag{S22}$$

and for a male of age $a$:

$$\gamma(m_\mathrm{a}) = \zeta\left(\sum_{x=a}^{\infty}\phi_\mathrm{f}(x)(1-\mu_\mathrm{f}(x,a))\right) + (1-\zeta)\left(\sum_{x=a}^{\infty}\phi_\mathrm{f}(x)\mu_\mathrm{m}(x,a)\right) = \zeta\gamma_\mathrm{f}(m_\mathrm{a}) + (1-\zeta)\gamma_\mathrm{m}(m_\mathrm{a}) \tag{S23}$$

Once again, we can calculate the expected time of a succcessful loop. First for a female of age $a$:

$$\begin{aligned} t_\bullet(f_\mathrm{a}) &= \left(\frac{\zeta\gamma_\mathrm{f}(f_\mathrm{a})}{\gamma(f_\mathrm{a})}\right)\left(\frac{\sum_{x=a}^{\infty}\phi_\mathrm{f}(x)\mu_\mathrm{f}(x,a)x}{\sum_{x=a}^{\infty}\phi_\mathrm{f}(x)\mu_\mathrm{f}(x,a)}\right) + \left(\frac{(1-\zeta)\gamma_\mathrm{m}(f_\mathrm{a})}{\gamma(f_\mathrm{a})}\right)\left(\frac{\sum_{x=a}^{\infty}\phi_\mathrm{m}(x)(1-\mu_\mathrm{m}(x,a))x}{\sum_{x=a}^{\infty}\phi_\mathrm{m}(x)(1-\mu_\mathrm{m}(x,a))}\right) \\ &= \left(\frac{\zeta\gamma_\mathrm{f}(f_\mathrm{a})}{\gamma(f_\mathrm{a})}\right)\left(\frac{T_\bullet^\mathrm{f}(f_\mathrm{a})}{\gamma_\mathrm{f}(f_\mathrm{a})}\right) + \left(\frac{(1-\zeta)\gamma_\mathrm{m}(f_\mathrm{a})}{\gamma(f_\mathrm{a})}\right)\left(\frac{T_\bullet^\mathrm{m}(f_\mathrm{a})}{\gamma_\mathrm{m}(f_\mathrm{a})}\right) \end{aligned} \tag{S24}$$

And for a male of age $a$:

$$\begin{aligned} t_\bullet(m_\mathrm{a}) &= \left(\frac{\zeta\gamma_\mathrm{f}(f_\mathrm{a})}{\gamma(f_\mathrm{a})}\right)\left(\frac{\sum_{x=a}^{\infty}\phi_\mathrm{f}(x)(1-\mu_\mathrm{f}(x,a))x}{\sum_{x=a}^{\infty}\phi_\mathrm{f}(x)(1-\mu_\mathrm{f}(x,a))}\right) + \left(\frac{(1-\zeta)\gamma_\mathrm{m}(f_\mathrm{a})}{\gamma(f_\mathrm{a})}\right)\left(\frac{\sum_{x=a}^{\infty}\phi_\mathrm{m}(x)\mu_\mathrm{m}(x,a)x}{\sum_{x=a}^{\infty}\phi_\mathrm{m}(x)\mu_\mathrm{m}(x,a)}\right) \\ &= \left(\frac{\zeta\gamma_\mathrm{f}(f_\mathrm{a})}{\gamma(f_\mathrm{a})}\right)\left(\frac{T_\bullet^\mathrm{f}(f_\mathrm{a})}{\gamma_\mathrm{f}(f_\mathrm{a})}\right) + \left(\frac{(1-\zeta)\gamma_\mathrm{m}(f_\mathrm{a})}{\gamma(f_\mathrm{a})}\right)\left(\frac{T_\bullet^\mathrm{m}(f_\mathrm{a})}{\gamma_\mathrm{m}(f_\mathrm{a})}\right) \end{aligned} \tag{S25}$$

We can also write out the expected time of an unsuccessful loop. First, for a female of age $a$:

$$\begin{aligned} t_\circ(f_\mathrm{a}) &= \left(\frac{\zeta(1-\gamma_\mathrm{f}(f_\mathrm{a}))}{1-\gamma(f_\mathrm{a})}\right)\left(\frac{\sum_{x=a}^{a-1}\phi_\mathrm{f}(x)x + \sum_{x=a}^{\infty}\phi_\mathrm{f}(x)(1-\mu_\mathrm{f}(x,a))x}{\sum_{x=a}^{a-1}\phi_\mathrm{f}(x) + \sum_{x=a}^{\infty}\phi_\mathrm{f}(x)(1-\mu_\mathrm{f}(x,a))}\right) \\ &+ \left(\frac{(1-\zeta)(1-\gamma_\mathrm{m}(f_\mathrm{a}))}{1-\gamma(f_\mathrm{a})}\right)\left(\frac{\sum_{x=a}^{a-1}\phi_\mathrm{m}(x)x + \sum_{x=a}^{\infty}\phi_\mathrm{m}(x)\mu_\mathrm{m}(x,a)x}{\sum_{x=a}^{a-1}\phi_\mathrm{m}(x) + \sum_{x=a}^{\infty}\phi_\mathrm{m}(x)\mu_\mathrm{m}(x,a)}\right) \\ &= \left(\frac{\zeta(1-\gamma_\mathrm{f}(f_\mathrm{a}))}{1-\gamma(f_\mathrm{a})}\right)\left(\frac{T_\circ^\mathrm{f}(f_\mathrm{a})}{1-\gamma_\mathrm{f}(f_\mathrm{a})}\right) + \left(\frac{(1-\zeta)(1-\gamma_\mathrm{m}(f_\mathrm{a}))}{1-\gamma(f_\mathrm{a})}\right)\left(\frac{T_\circ^\mathrm{m}(f_\mathrm{a})}{1-\gamma_\mathrm{m}(f_\mathrm{a})}\right) \end{aligned} \tag{S26}$$

And for a male of age $a$:

$$\begin{aligned} t_\circ(m_\mathrm{a}) &= \left(\frac{\zeta(1-\gamma_\mathrm{f}(m_\mathrm{a}))}{1-\gamma(m_\mathrm{a})}\right)\left(\frac{\sum_{x=a}^{a-1}\phi_\mathrm{f}(x)x + \sum_{x=a}^{\infty}\phi_\mathrm{f}(x)\mu_\mathrm{f}(x,a)x}{\sum_{x=a}^{a-1}\phi_\mathrm{f}(x) + \sum_{x=a}^{\infty}\phi_\mathrm{f}(x)\mu_\mathrm{f}(x,a)}\right) \\ &+ \left(\frac{(1-\zeta)(1-\gamma_\mathrm{m}(m_\mathrm{a}))}{1-\gamma(m_\mathrm{a})}\right)\left(\frac{\sum_{x=a}^{a-1}\phi_\mathrm{m}(x)x + \sum_{x=a}^{\infty}\phi_\mathrm{m}(x)(1-\mu_\mathrm{m}(x,a))x}{\sum_{x=a}^{a-1}\phi_\mathrm{m}(x) + \sum_{x=a}^{\infty}\phi_\mathrm{m}(x)(1-\mu_\mathrm{m}(x,a))}\right) \\ &= \left(\frac{\zeta(1-\gamma_\mathrm{f}(m_\mathrm{a}))}{1-\gamma(m_\mathrm{a})}\right)\left(\frac{T_\circ^\mathrm{f}(m_\mathrm{a})}{1-\gamma_\mathrm{f}(m_\mathrm{a})}\right) + \left(\frac{(1-\zeta)(1-\gamma_\mathrm{m}(m_\mathrm{a}))}{1-\gamma(m_\mathrm{a})}\right)\left(\frac{T_\circ^\mathrm{m}(m_\mathrm{a})}{1-\gamma_\mathrm{m}(m_\mathrm{a})}\right) \end{aligned} \tag{S27}$$

And again we can put these together to recover the mean return times. First of a female of age $a$:

$$\begin{aligned}\mathscr{R}(f_\mathrm{a}) &= t_\bullet(f_\mathrm{a}) + \left(\frac{1-\gamma(f_\mathrm{a})}{\gamma(f_\mathrm{a})}\right) t_\circ(f_\mathrm{a}) \\ &= \frac{\zeta T_\mathrm{f} + (1-\zeta) T_\mathrm{m}}{\gamma(f_\mathrm{a})} = \frac{\tilde{T}}{\gamma(f_\mathrm{a})}\end{aligned} \tag{S28}$$

And a male of age $a$:

$$\begin{aligned}\mathscr{R}(m_\mathrm{a}) &= t_\bullet(m_\mathrm{a}) + \left(\frac{1-\gamma(m_\mathrm{a})}{\gamma(m_\mathrm{a})}\right) t_\circ(m_\mathrm{a}) \\ &= \frac{\zeta T_\mathrm{f} + (1-\zeta) T_\mathrm{m}}{\gamma(m_\mathrm{a})} = \frac{\tilde{T}}{\gamma(m_\mathrm{a})}\end{aligned} \tag{S29}$$

Once again, we can convert these into class reproductive values:

$$c(f_\mathrm{a}) = \gamma(f_\mathrm{a}) / \tilde{T} \tag{S30}$$

And:

$$c(m_\mathrm{a}) = \gamma(m_\mathrm{a}) / \tilde{T} \tag{S31}$$

We can also sum across some of these different classes. Summing across individuals of age 1:

$$c(1) = c(f_1) + c(m_1) = \frac{\gamma(f_1) + \gamma(m_1)}{\tilde{T}} = \frac{\zeta \sum_{x=1}^{\infty} \phi_\mathrm{f}(x) + (1-\zeta) \sum_{x=1}^{\infty} \phi_\mathrm{m}(x)}{\tilde{T}} = \frac{1}{\tilde{T}} \tag{S32}$$

Recovering the result for simultaneous hermaphrodites. And for other ages:

$$c(a) = c(f_\mathrm{a}) + c(m_\mathrm{a}) = \frac{\gamma(f_\mathrm{a}) + \gamma(m_\mathrm{a})}{\tilde{T}} = \frac{\zeta \sum_{x=a}^{\infty} \phi_\mathrm{f}(x) + (1-\zeta) \sum_{x=a}^{\infty} \phi_\mathrm{m}(x)}{\tilde{T}} \tag{S33}$$

Which again is equivalent to expression for simultaneous hermaphrodites. If we sum across all individuals of the same sex, then for females:

$$c(f) = \sum_{a}^{\infty} c(f_\mathrm{a}) = \frac{1}{\tilde{T}} \sum_{a} \gamma(f_\mathrm{a}) = \gamma(f_\mathrm{a}) = \frac{1}{\tilde{T}} \sum_{a} \left( \zeta \left( \sum_{x=a}^{\infty} \phi_\mathrm{f}(x) \mu_\mathrm{f}(x,a) \right) + (1-\zeta) \left( \sum_{x=a}^{\infty} \phi_\mathrm{f}(x)(1-\mu_\mathrm{m}(x,a)) \right) \right) \tag{S34}$$

Which we may rewrite this in a slightly different way:

$$\begin{aligned}c(f) &= \frac{1}{\tilde{T}} \left( \zeta \sum_{a} \sum_{x=a}^{\infty} \left( \phi_\mathrm{f}(x) \mu_\mathrm{f}(x,a) \right) + (1-\zeta) \sum_{a} \sum_{x=a}^{\infty} \left( \phi_\mathrm{m}(x)(1-\mu_\mathrm{f}(x,a)) \right) \right) \\ &= \frac{1}{\tilde{T}} \left( \zeta \sum_{x=1}^{\infty} \left( \phi_\mathrm{f}(x) \rho_\mathrm{f}(x) x \right) + \zeta \sum_{x=1}^{\infty} \left( \phi_\mathrm{m}(x)(1-\rho_\mathrm{m}(x)) x \right) \right) \\ &= \frac{1}{\tilde{T}} \left( \zeta T_\mathrm{ff} + (1-\zeta) T_\mathrm{mf} \right)\end{aligned} \tag{S35}$$

Where: $\rho_\mathrm{f}(x)$ is the mean proportion of an age $x$ female's life that she has spent as a female, $\rho_\mathrm{m}(x)$ is the mean proportion of an age $x$ male's life that he has spent as a male, $T_\mathrm{ff}$ is the mean time that a mother of a newborn has spent as a female, and $T_\mathrm{mf}$ is the mean time that a father of a newborn has spent as a male. Similarly for males:

$$\begin{aligned}c(m) &= \sum_{a}^{\infty} c(m_\mathrm{a}) \\ &= \frac{1}{\tilde{T}} \left( \zeta \sum_{x=1}^{\infty} \left( \phi_\mathrm{f}(x)(1-\rho_\mathrm{f}(x)) x \right) + \zeta \sum_{x=1}^{\infty} \left( \phi_\mathrm{m}(x) \rho_\mathrm{m}(x) x \right) \right) \\ &= \frac{1}{\tilde{T}} \left( \zeta T_\mathrm{fm} + (1-\zeta) T_\mathrm{mm} \right)\end{aligned} \tag{S36}$$

The ratio of these two is presented in the main text (Equation 1). As before, we may partition our class reproductive values into survival and reproduction. Through reproduction:

$$c_{\mathrm{r}}(f_{\mathrm{a}}) = \zeta\phi_{\mathrm{f}}(a)c(1) = \zeta\frac{\phi_{\mathrm{f}}(a)}{\tilde{T}} \tag{S37}$$

And:

$$c_{\mathrm{r}}(m_{\mathrm{a}}) = (1-\zeta)\phi_{\mathrm{m}}(a)c(1) = (1-\zeta)\frac{\phi_{\mathrm{m}}(a)}{\tilde{T}} \tag{S38}$$

And through survival:

$$\begin{aligned} c_s(f_{\mathrm{a}}) =& \mu_{\mathrm{f}}(a+1,a)c(f_{\mathrm{a+1}}) + \big(1-\mu_{\mathrm{m}}(a+1,a)\big)\,c(m_{\mathrm{a+1}}) \\ =& \frac{1}{\tilde{T}}\left(\mu_{\mathrm{f}}(a+1,a)\gamma(f_{\mathrm{a+1}}) + \big(1-\mu_{\mathrm{m}}(a+1,a)\big)\gamma(m_{\mathrm{a+1}})\right) \end{aligned} \tag{S39}$$

$$\begin{aligned} c_s(m_{\mathrm{a}}) =& \big(1-\mu_{\mathrm{f}}(a+1,a)\big)\,c(f_{\mathrm{a+1}}) + \mu_{\mathrm{m}}(a+1,a)c(m_{\mathrm{a+1}}) \\ =& \frac{1}{\tilde{T}}\left(\big(1-\mu_{\mathrm{f}}(a+1,a)\big)\gamma(f_{\mathrm{a+1}}) + \mu_{\mathrm{m}}(a+1,a)\gamma(m_{\mathrm{a+1}})\right) \end{aligned} \tag{S40}$$

#### 2.5 Sexual reproduction: gonochorism

Now, we consider gonochorism (or dioecy). Whilst one form of gonochorism may be recovered from the previous section by simply setting $\mu_{\mathrm{f}} = \mu_{\mathrm{m}} = 1$, we now extend this by allowing for scenarios where there are different patterns of ancestry between males and females, for example asymmetric genetic systems such as haplodiploidy. We introduce the following additional notation: $\zeta_{\mathrm{f}}$ is the probability that a gene copy in a female came from a female and $\zeta_{\mathrm{m}}$ is the probability that a gene copy in a male came from a male.

With this, we may write the return time for a gene copy in a newborn female as:

$$\mathscr{R}(f_1) = \zeta_{\mathrm{f}}T_{\mathrm{f}} + (1-\zeta_{\mathrm{f}})\left(T_{\mathrm{m}} + \left(\frac{1}{1-\zeta_{\mathrm{m}}} - 1\right)T_{\mathrm{m}} + T_{\mathrm{f}}\right) = T_{\mathrm{f}} + \left(\frac{1-\zeta_{\mathrm{f}}}{1-\zeta_{\mathrm{m}}}\right)T_{\mathrm{m}} \tag{S41}$$

And for a newborn male is:

$$\mathscr{R}(m_1) = T_{\mathrm{m}} + \left(\frac{1-\zeta_{\mathrm{m}}}{1-\zeta_{\mathrm{f}}}\right)T_{\mathrm{f}} \tag{S42}$$

And so the class reproductive value of newborns in total is:

$$c(1) = c(f_1) + c(m_1) = \frac{1}{\mathscr{R}(f_1)} + \frac{1}{\mathscr{R}(m_1)} = \frac{(1-\zeta_{\mathrm{f}}) + (1-\zeta_{\mathrm{m}})}{(1-\zeta_{\mathrm{f}})T_{\mathrm{f}} + (1-\zeta_{\mathrm{m}})T_{\mathrm{m}}} = \frac{1}{\tilde{T}} \tag{S43}$$

Where:

$$\tilde{T} = T_{\mathrm{f}}\left(\frac{1-\zeta_{\mathrm{m}}}{(1-\zeta_{\mathrm{f}}) + (1-\zeta_{\mathrm{m}})}\right) + T_{\mathrm{m}}\left(\frac{1-\zeta_{\mathrm{f}}}{(1-\zeta_{\mathrm{f}}) + (1-\zeta_{\mathrm{m}})}\right) \tag{S44}$$

And the ratio of these two quantities is:

$$\frac{c(f_1)}{c(m_1)} = \frac{1-\zeta_{\mathrm{m}}}{1-\zeta_{\mathrm{f}}} \tag{S45}$$

This recovers the result of Hitchcock and Gardner (2020), through a slightly different approach. We can, as before, consider the mean return time for a gene of a particular age and sex. First, let's consider a female of age $a$. It takes her $a$ time units to return to the newborn state. Let's define $\mathscr{R}(x \to y)$ as the mean time from state $x$

to state $y$. The mean time from the female newborn state to a female of age $a$ is:

$$\begin{aligned}\mathscr{R}(f_1 \to f_a) = &\left(\frac{\big(1-\gamma_f(a)\big)\zeta_f}{\gamma_f(a)\zeta_f + (1-\zeta_f)}\right)\left(\frac{T_{f\circ}}{1-\gamma_f(a)}\right)\\ &+\left(\frac{1-\zeta_f}{\gamma_f(a)\zeta_f + (1-\zeta_f)}\right)\big(T_m + \mathscr{R}(m_1 \to f_a)\big)\\ &+\left(\frac{\gamma_f(a)\zeta_f}{\gamma_f(a)\zeta_f + (1-\zeta_f)}\right)\left(\frac{T_{f\bullet}}{\gamma_f(a)} - a\right)\end{aligned} \tag{S46}$$

Where:

$$\gamma_f(a) = \sum_{x=a}^{\infty} \phi_f(x) \tag{S47}$$

And the mean time from the newborn male state:

$$\begin{aligned}\mathscr{R}(m_1 \to f_a) = &\left(\frac{\zeta_m}{1-\zeta_m}\right)T_m\\ &+\left(\frac{(1-\zeta_m)\big(1-\gamma_f(a)\big)}{1-\zeta_f}\right)\left(\frac{T_{f\circ}}{1-\gamma_f(a)} + \mathscr{R}(f_1 \to f_a)\right)\\ &+\left(\frac{(1-\zeta_m)\gamma_f(a)}{1-\zeta_f}\right)\left(\frac{T_{f\bullet}}{\gamma_f}(a) - a\right)\end{aligned} \tag{S48}$$

We may then solve these simultaneous equations for $\mathscr{R}(f_1 \to f_a)$. Combining and simplifying, we can then write out our return time for $f_a$ as:

$$\mathscr{R}(f_a) = a + \mathscr{R}(f_1 \to f_a) = \frac{\tilde{T}}{\left(\frac{1-\zeta_m}{(1-\zeta_f)+(1-\zeta_m)}\right)\gamma_f(a)} \tag{S49}$$

And similarly, we can do the same procedure to calculate the mean return time for a male of age $a$.

$$\mathscr{R}(m_a) = a + \mathscr{R}(m_1 \to m_a) = \frac{\tilde{T}}{\left(\frac{1-\zeta_f}{(1-\zeta_f)+(1-\zeta_m)}\right)\gamma_m(a)} \tag{S50}$$

Again, these can be be inverted to get the class reproductive values:

$$c(f_a) = \frac{1}{\mathscr{R}(f_a)} = \left(\frac{1-\zeta_m}{(1-\zeta_f)+(1-\zeta_m)}\right)\frac{\gamma_f(a)}{\tilde{T}} \tag{S51}$$

$$c(m_a) = \frac{1}{\mathscr{R}(m_a)} = \left(\frac{1-\zeta_f}{(1-\zeta_f)+(1-\zeta_m)}\right)\frac{\gamma_m(a)}{\tilde{T}} \tag{S52}$$

Where the left side of each expression can be viewed as weighting how the genetics shapes the distribution of reproductive values, and the left portion the demography. Once again, we may further partition these expressions into the contributions through survival and reproduction. For females, reproduction through daughters $c^r_{f\to f}(a) = \zeta_f\phi_f(a)c(f_1)$ and through sons $c^r_{f\to m}(a) = (1-\zeta_m)\phi_f(a)c(m_1)$, and through survival $c^s_{f\to f}(a) = c(f_{a+1})$. Similarly for males, class reproductive value through daughters:$c^r_{m\to f}(a) = (1-\zeta_f)\phi_m(a)c(m_1)$ and through sons $c^r_{m\to m}(a) = \zeta_m\phi_f(a)c(m_1)$, and through survival $c^s_{m\to m}(a) = c(m_{a+1})$.

### 3 Population genetics model

For the purpose of concreteness and illustration, we analyse a simple model that incorporates two possible age-classes ($a \in \{1,2\}$) and two possible sex-classes ($k \in \{f, m\}$), as depicted in Figure S1. Within this structure, we allow for a variety of sex change systems by describing the following parameters: $\alpha$ is the fraction of age-2 who females came from age-1 females in the previous generation, $\beta$ is the fraction of age-2 males who came from age-1 males in the previous generation, $\mu$ is the fraction of newborns who have age-1 mothers, and $\nu$ is the fraction of newborns who have age-1 fathers. Thus, simple gonochorism can be recovered by setting $\alpha = \beta = 1$, simple protandry can be recovered by setting $\alpha = 0$ and $\beta = 1$, and simple protogyny can be recovered by setting $\alpha = 1$ and $\beta = 0$.

Using this model we may compute the various invasion conditions for different types of sexually antagonistic trade-offs in terms of the above parameters. We first present results for arbitrary strength of selection, and then derive weak selection approximations from these. We assume that there is no population structure or assortative mating. We rearrange our weak selection approximations into the forms of "potentials for feminisation", and these results can be seen in Table 2 of the main text.

The potential for feminisation for various combinations of these trade-offs for both autosomal and cytoplasmic genes are given in Table 2 of the main text. We provide numerical illustrations of the resulting potential for polymorphism for a subset of these scenarios, under the assumption of weak selection, for a range of different sexually antagonistic trade-offs in Figures 2 and 3.

#### 3.1 Reproductive values

For the life cycle described above and in Figure S1, we compute the reproductive values. To do this we may calculate the left ($\vec{u}$) and right ($\vec{v}$) eigenvectors associated with the dominant eigenvalue ($\lambda$) of our backwards transition matrix $T$.

$$\lambda \vec{u} = \vec{u} T \tag{S53}$$

$$\lambda \vec{v} = T \vec{v} \tag{S54}$$

Where the normalised vector of class reproductive values is:

$$\vec{c} = \frac{\vec{u} \times \vec{v}}{\vec{u} \cdot \vec{v}} \tag{S55}$$

This approach also would work if we were working directly with a projection matrix (Caswell, 2000). Which gives us the following class reproductive values:

$$\vec{c} = \begin{pmatrix} c_{\mathrm{f}_1} & c_{\mathrm{f}_2} & c_{\mathrm{m}_1} & c_{\mathrm{m}_2} \end{pmatrix} \tag{S56}$$

Where:

$$\tilde{T} = \zeta\left(\mu + 2(1-\mu)\right) + (1-\zeta)\left(\nu + 2(1-\nu)\right) \tag{S57a}$$

$$c_{\mathrm{f}_1} = \frac{\zeta\mu + \alpha\zeta(1-\mu) + (1-\beta)(1-\zeta)(1-\nu)}{\tilde{T}} \tag{S57b}$$

$$c_{\mathrm{f}_2} = \frac{\zeta(1-\mu)}{\tilde{T}} \tag{S57c}$$

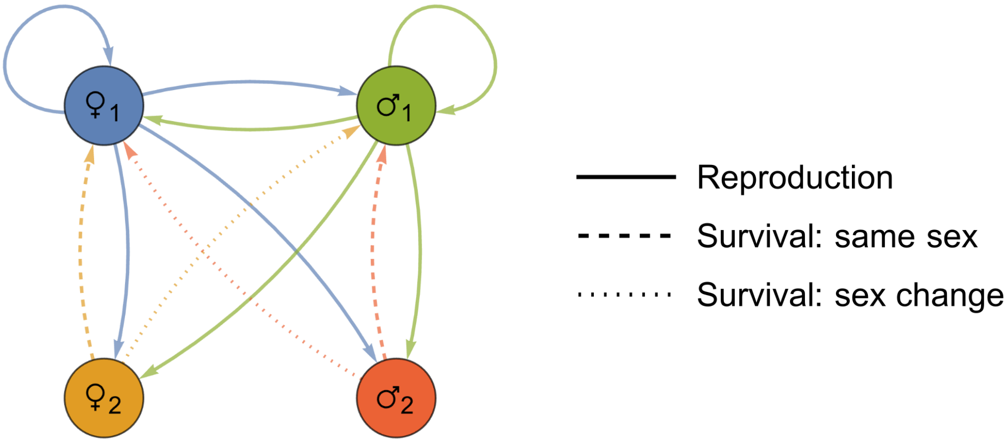


$$\begin{pmatrix} \phi_{♀_1\to♀_1} & \phi_{♀_1\to♀_2} & \phi_{♀_1\to♂_1} & \phi_{♀_1\to♂_2} \\ \phi_{♀_2\to♀_1} & \phi_{♀_2\to♀_2} & \phi_{♀_2\to♂_1} & \phi_{♀_2\to♂_2} \\ \phi_{♂_1\to♀_1} & \phi_{♂_1\to♀_2} & \phi_{♂_1\to♂_1} & \phi_{♂_1\to♂_2} \\ \phi_{♂_2\to♀_1} & \phi_{♂_2\to♀_2} & \phi_{♂_2\to♂_1} & \phi_{♂_2\to♂_2} \end{pmatrix} = \begin{pmatrix} \zeta\,\mu & \zeta\,(1-\mu) & (1-\zeta)\,v & (1-\zeta)\,(1-v) \\ \alpha & 0 & 1-\alpha & 0 \\ \zeta\,\mu & \zeta\,(1-\mu) & (1-\zeta)\,v & (1-\zeta)\,(1-v) \\ 1-\beta & 0 & \beta & 0 \end{pmatrix}$$

Figure S1: **Description of the life-cycle analysed in the population genetic model.** A) Graph demonstrating the flow of genes between different age and sex classes when traced back through time. The type of line indicates the type of transition that can occur (reproduction/survival/sex change). B) The associated transition matrix.

$$c_{m_1} = \frac{(1-\zeta)\nu + (1-\alpha)\zeta(1-\mu) + (1-\nu)\beta(1-\zeta)}{\tilde{T}} \tag{S57d}$$

$$c_{m_2} = \frac{(1-\zeta)(1-\nu)}{\tilde{T}} \tag{S57e}$$

We can then partition these reproductive values once more into the different routes through the life-cycle:

$$E = \begin{pmatrix} c_{f_1\to f_1} & c_{f_1\to f_2} & c_{f_1\to m_1} & c_{f_1\to m_2} \\ c_{f_2\to f_1} & c_{f_2\to f_2} & c_{f_2\to m_1} & c_{f_2\to m_2} \\ c_{m_1\to f_1} & c_{m_1\to f_2} & c_{m_1\to m_1} & c_{m_1\to m_2} \\ c_{m_2\to f_1} & c_{m_2\to f_2} & c_{m_2\to m_1} & c_{m_2\to m_2} \end{pmatrix} \tag{S58}$$

We can then sum the different types of routes through the life cycle which we will use later:

$$c_f^r = c_{f_1\to f_1} + c_{f_1\to m_1} + c_{f_2\to f_1} + c_{f_2\to m_1} = \frac{\zeta}{\tilde{T}} \tag{S59}$$

$$c_m^r = c_{m_1\to f_1} + c_{m_1\to m_1} + c_{m_2\to f_1} + c_{m_2\to m_1} = \frac{1-\zeta}{\tilde{T}} \tag{S60}$$

$$c_f^s = c_{f_1\to f_2} + c_{f_1\to m_2} = \frac{\alpha\zeta(1-\mu) + (1-\beta)(1-\zeta)(1-\nu)}{\tilde{T}} \tag{S61}$$

$$c_m^s = c_{m_1\to f_2} + c_{m_1\to m_2} = \frac{(1-\alpha)\zeta(1-\mu) + \beta(1-\zeta)(1-\nu)}{\tilde{T}} \tag{S62}$$

#### 3.2 Recursion equations

We first write out recursion equations describing the frequency of our different genotypes. As there are 4 states, then for the haploid case, we require $4\times(2-1) = 4$ equations, and for our diploid system we require $4\times(3-1) = 2$ equations. For the haploid system we consider potential asymmetries in transmission genetics, for the diploid system we assume that transmission genetics are symmetric between males and females ($\zeta = 1/2$).

##### 3.2.1 Haploid

Age 1 individuals

$$p'_{\{f_1,1\}} = p'_{\{m_1,1\}} = \zeta\left(\mu p_{\{f_1,1\}}\frac{\omega_{\{f_1,1\}}}{\overline{\omega}_{\{f_1\}}} + (1-\mu)p_{\{f_2,1\}}\frac{\omega_{\{f_2,1\}}}{\overline{\omega}_{\{f_2\}}}\right) + (1-\zeta)\left(\nu p_{\{m_1,1\}}\frac{\omega_{\{m_1,1\}}}{\overline{\omega}_{\{m_1\}}} + (1-\nu)p_{\{m_2,1\}}\frac{\omega_{\{m_2,1\}}}{\overline{\omega}_{\{m_2\}}}\right) \tag{S63}$$

Where:

$$\overline{\omega}_{\{f_1\}} = \left(1 - p_{\{f_1,1\}}\right)\omega_{\{f_1,0\}} + p_{\{f_1,1\}}\omega_{\{f_1,1\}} \tag{S64a}$$

$$\overline{\omega}_{\{f_2\}} = \left(1 - p_{\{f_2,1\}}\right)\omega_{\{f_2,0\}} + p_{\{f_2,1\}}\omega_{\{f_2,1\}} \tag{S64b}$$

$$\overline{\omega}_{\{m_1\}} = \left(1 - p_{\{m_1,1\}}\right)\omega_{\{m_1,0\}} + p_{\{m_1,1\}}\omega_{\{m_1,1\}} \tag{S64c}$$

$$\overline{\omega}_{\{m_2\}} = \left(1 - p_{\{m_2,1\}}\right)\omega_{\{m_2,0\}} + p_{\{m_2,1\}}\omega_{\{m_2,1\}} \tag{S64d}$$

Age 2 individuals

$$p'_{\{f_2,1\}} = \alpha\left(p_{\{f_1,1\}}\frac{\Omega_{\{f_1,1\}}}{\overline{\Omega}_{\{f_1\}}}\right) + (1-\alpha)\left(p_{\{m_1,1\}}\frac{\Omega_{\{m_1,1\}}}{\overline{\Omega}_{\{m_1\}}}\right) \tag{S65}$$

$$p'_{\{\mathrm{m}_2,1\}} = (1-\beta)\left(p_{\{\mathrm{f}_1,1\}}\frac{\Omega_{\{\mathrm{f}_1,1\}}}{\overline{\Omega}_{\{\mathrm{f}_1\}}}\right) + \beta\left(p_{\{\mathrm{m}_1,1\}}\frac{\Omega_{\{\mathrm{m}_1,1\}}}{\overline{\Omega}_{\{\mathrm{m}_1\}}}\right) \tag{S66}$$

Where:

$$\overline{\Omega}_{\{\mathrm{f}_1\}} = \left(1-p_{\{\mathrm{m}_1,1\}}\right)\Omega_{\{\mathrm{f}_1,0\}} + p_{\{\mathrm{f}_1,1\}}\Omega_{\{\mathrm{f}_1,1\}} \tag{S67a}$$

$$\overline{\Omega}_{\{\mathrm{m}_1\}} = \left(1-p_{\{\mathrm{m}_1,1\}}\right)\Omega_{\{\mathrm{m}_1,0\}} + p_{\{\mathrm{m}_1,1\}}\Omega_{\{\mathrm{m}_1,1\}} \tag{S67b}$$

##### 3.2.2 Diploid

Age 1 individuals:

$$\begin{aligned} p'_{\{\mathrm{f}_1,01\}} = p'_{\{\mathrm{m}_1,01\}} = &\\ &\left(\mu\left(\frac{p_{\{\mathrm{f}_1,00\}}\omega_{\{\mathrm{f}_1,00\}} + \frac{1}{2}p_{\{\mathrm{f}_1,01\}}\omega_{\{\mathrm{f}_1,01\}} + \frac{1}{2}p_{\{\mathrm{f}_1,10\}}\omega_{\{\mathrm{f}_1,10\}}}{\overline{\omega}_{\{\mathrm{f}_1\}}}\right) + \right.\\ &\left.(1-\mu)\left(\frac{p_{\{\mathrm{f}_2,00\}}\omega_{\{\mathrm{f}_2,00\}} + \frac{1}{2}p_{\{\mathrm{f}_2,01\}}\omega_{\{\mathrm{f}_2,01\}} + \frac{1}{2}p_{\{\mathrm{f}_2,10\}}\omega_{\{\mathrm{f}_2,10\}}}{\overline{\omega}_{\{\mathrm{f}_2\}}}\right)\right) \times\\ &\left(\nu\left(\frac{\frac{1}{2}p_{\{\mathrm{m}_1,01\}}\omega_{\{\mathrm{m}_1,01\}} + \frac{1}{2}p_{\{\mathrm{m}_1,10\}}\omega_{\{\mathrm{m}_1,10\}} + p_{\{\mathrm{m}_1,11\}}\omega_{\{\mathrm{m}_1,11\}}}{\overline{\omega}_{\{\mathrm{m}_1\}}}\right) + \right.\\ &\left.(1-\nu)\left(\frac{\frac{1}{2}p_{\{\mathrm{m}_2,01\}}\omega_{\{\mathrm{m}_2,01\}} + \frac{1}{2}p_{\{\mathrm{m}_2,10\}}\omega_{\{\mathrm{m}_2,10\}} + p_{\{\mathrm{m}_2,11\}}\omega_{\{\mathrm{m}_2,11\}}}{\overline{\omega}_{\{\mathrm{m}_2\}}}\right)\right) \end{aligned} \tag{S68}$$

$$\begin{aligned} p'_{\{\mathrm{f}_1,10\}} = p'_{\{\mathrm{m}_1,10\}} = &\\ &\left(\mu\left(\frac{\frac{1}{2}p_{\{\mathrm{f}_1,01\}}\omega_{\{\mathrm{f}_1,01\}} + \frac{1}{2}p_{\{\mathrm{f}_1,10\}}\omega_{\{\mathrm{f}_1,10\}} + p_{\{\mathrm{f}_1,11\}}\omega_{\{\mathrm{f}_1,11\}}}{\overline{\omega}_{\{\mathrm{f}_1\}}}\right) + \right.\\ &\left.(1-\mu)\left(\frac{\frac{1}{2}p_{\{\mathrm{f}_2,01\}}\omega_{\{\mathrm{f}_2,01\}} + \frac{1}{2}p_{\{\mathrm{f}_2,10\}}\omega_{\{\mathrm{f}_2,10\}} + p_{\{\mathrm{f}_2,11\}}\omega_{\{\mathrm{f}_2,11\}}}{\overline{\omega}_{\{\mathrm{f}_2\}}}\right)\right) \times\\ &\left(\nu\left(\frac{p_{\{\mathrm{m}_1,00\}}\omega_{\{\mathrm{m}_1,00\}} + \frac{1}{2}p_{\{\mathrm{m}_1,01\}}\omega_{\{\mathrm{m}_1,01\}} + \frac{1}{2}p_{\{\mathrm{m}_1,10\}}\omega_{\{\mathrm{m}_1,10\}}}{\overline{\omega}_{\{\mathrm{m}_1\}}}\right) + \right.\\ &\left.(1-\nu)\left(\frac{p_{\{\mathrm{m}_2,00\}}\omega_{\{\mathrm{m}_2,00\}} + \frac{1}{2}p_{\{\mathrm{m}_2,01\}}\omega_{\{\mathrm{m}_2,01\}} + \frac{1}{2}p_{\{\mathrm{m}_2,10\}}\omega_{\{\mathrm{m}_2,10\}}}{\overline{\omega}_{\{\mathrm{m}_2\}}}\right)\right) \end{aligned} \tag{S69}$$

$$\begin{aligned} p'_{\{\mathrm{f}_1,11\}} = p'_{\{\mathrm{m}_1,11\}} = &\\ &\left(\mu\left(\frac{\frac{1}{2}p_{\{\mathrm{f}_1,01\}}\omega_{\{\mathrm{f}_1,01\}} + \frac{1}{2}p_{\{\mathrm{f}_1,10\}}\omega_{\{\mathrm{f}_1,10\}} + p_{\{\mathrm{f}_1,11\}}\omega_{\{\mathrm{f}_1,11\}}}{\overline{\omega}_{\{\mathrm{f}_1\}}}\right) + \right.\\ &\left.(1-\mu)\left(\frac{\frac{1}{2}p_{\{\mathrm{f}_2,01\}}\omega_{\{\mathrm{f}_2,01\}} + \frac{1}{2}p_{\{\mathrm{f}_2,10\}}\omega_{\{\mathrm{f}_2,10\}} + p_{\{\mathrm{f}_2,11\}}\omega_{\{\mathrm{f}_2,11\}}}{\overline{\omega}_{\{\mathrm{f}_2\}}}\right)\right) \times\\ &\left(\nu\left(\frac{\frac{1}{2}p_{\{\mathrm{m}_1,01\}}\omega_{\{\mathrm{m}_1,01\}} + \frac{1}{2}p_{\{\mathrm{m}_1,10\}}\omega_{\{\mathrm{m}_1,10\}} + p_{\{\mathrm{m}_1,11\}}\omega_{\{\mathrm{m}_1,11\}}}{\overline{\omega}_{\{\mathrm{m}_1\}}}\right) + \right.\\ &\left.(1-\nu)\left(\frac{\frac{1}{2}p_{\{\mathrm{m}_2,01\}}\omega_{\{\mathrm{m}_2,01\}} + \frac{1}{2}p_{\{\mathrm{m}_2,10\}}\omega_{\{\mathrm{m}_2,10\}} + p_{\{\mathrm{m}_2,11\}}\omega_{\{\mathrm{m}_2,11\}}}{\overline{\omega}_{\{\mathrm{m}_2\}}}\right)\right) \end{aligned} \tag{S70}$$

Where:

$$\overline{\omega}_{\{\mathrm{f}_1\}} = p_{\{\mathrm{f}_1,00\}}\omega_{\{\mathrm{f}_1,00\}} + p_{\{\mathrm{f}_1,01\}}\omega_{\{\mathrm{f}_1,01\}} + p_{\{\mathrm{f}_1,10\}}\omega_{\{\mathrm{f}_1,10\}} + p_{\{\mathrm{f}_1,11\}}\omega_{\{\mathrm{f}_1,11\}} \tag{S71a}$$

$$\overline{\omega}_{\{f_2\}} = p_{\{f_2,00\}}\omega_{\{f_2,00\}} + p_{\{f_2,01\}}\omega_{\{f_2,01\}} + p_{\{f_2,10\}}\omega_{\{f_2,10\}} + p_{\{f_2,11\}}\omega_{\{f_2,11\}} \tag{S71b}$$

$$\overline{\omega}_{\{f_1\}} = p_{\{m_1,00\}}\omega_{\{m_1,00\}} + p_{\{m_1,01\}}\omega_{\{m_1,01\}} + p_{\{m_1,10\}}\omega_{\{m_1,10\}} + p_{\{m_1,11\}}\omega_{\{m_1,11\}} \tag{S71c}$$

$$\overline{\omega}_{\{f_2\}} = p_{\{m_2,00\}}\omega_{\{m_2,00\}} + p_{\{m_2,01\}}\omega_{\{m_2,01\}} + p_{\{m_2,10\}}\omega_{\{m_2,10\}} + p_{\{m_2,11\}}\omega_{\{m_2,11\}} \tag{S71d}$$

Age 2 individuals:

$$p'_{\{f_2,01\}} = \alpha\left(\frac{p_{\{f_1,01\}}\Omega_{\{f_1,01\}}}{\overline{\Omega}_{\{f_1\}}}\right) + (1-\alpha)\left(\frac{p_{\{m_1,01\}}\Omega_{\{m_1,01\}}}{\overline{\Omega}_{\{m_1\}}}\right) \tag{S72}$$

$$p'_{\{f_2,10\}} = \alpha\left(\frac{p_{\{f_1,10\}}\Omega_{\{f_1,10\}}}{\overline{\Omega}_{\{f_1\}}}\right) + (1-\alpha)\left(\frac{p_{\{m_1,10\}}\Omega_{\{m_1,10\}}}{\overline{\Omega}_{\{m_1\}}}\right) \tag{S73}$$

$$p'_{\{f_2,11\}} = \alpha\left(\frac{p_{\{f_1,11\}}\Omega_{\{f_1,11\}}}{\overline{\Omega}_{\{f_1\}}}\right) + (1-\alpha)\left(\frac{p_{\{m_1,11\}}\Omega_{\{m_1,11\}}}{\overline{\Omega}_{\{m_1\}}}\right) \tag{S74}$$

$$p'_{\{m_2,01\}} = (1-\beta)\left(\frac{p_{\{f_1,01\}}\Omega_{\{f_1,01\}}}{\overline{\Omega}_{\{f_1\}}}\right) + \beta\left(\frac{p_{\{m_1,01\}}\Omega_{\{m_1,01\}}}{\overline{\Omega}_{\{m_1\}}}\right) \tag{S75}$$

$$p'_{\{m_2,10\}} = (1-\beta)\left(\frac{p_{\{f_1,10\}}\Omega_{\{f_1,10\}}}{\overline{\Omega}_{\{f_1\}}}\right) + \beta\left(\frac{p_{\{m_1,10\}}\Omega_{\{m_1,10\}}}{\overline{\Omega}_{\{m_1\}}}\right) \tag{S76}$$

$$p'_{\{f_2,11\}} = (1-\beta)\left(\frac{p_{\{f_1,11\}}\Omega_{\{f_1,11\}}}{\overline{\Omega}_{\{f_1\}}}\right) + \beta\left(\frac{p_{\{m_1,11\}}\Omega_{\{m_1,11\}}}{\overline{\Omega}_{\{m_1\}}}\right) \tag{S77}$$

Where:

$$\overline{\Omega}_{\{f_1\}} = p_{\{f_1,00\}}\Omega_{\{f_1,00\}} + p_{\{f_1,01\}}\Omega_{\{f_1,01\}} + p_{\{f_1,10\}}\Omega_{\{f_1,10\}} + p_{\{f_1,11\}}\Omega_{\{f_1,11\}} \tag{S78a}$$

$$\overline{\Omega}_{\{m_1\}} = p_{\{m_1,00\}}\Omega_{\{m_1,00\}} + p_{\{m_1,01\}}\Omega_{\{m_1,01\}} + p_{\{m_1,10\}}\Omega_{\{m_1,10\}} + p_{\{m_1,11\}}\Omega_{\{m_1,11\}} \tag{S78b}$$

##### 3.3 Jacobians and eigenvalues

Using these recursion equations, we can then ask when the mutant allele will be able to invade from rarity. We consider the stability of the two trivial equilibria $p = 0$ and $p = 1$. To determine their stability, we first write out the Jacobian matrix **J**, analysed when the mutant (or resident) allele are vanishingly rare in the population (Otto and Day, 2011), i.e. a local linear stability analysis. Each entry of the matrix is given by:

$$\mathbf{J}_{i,j} = \left.\frac{\partial p'_i}{\partial p'_j}\right|_{p_*=0} \tag{S79}$$

If there exists at least one eigenvalue with real part strictly greater than zero, then the allele will be able to invade from rarity.

###### 3.3.1 Haploid

For the haploid system, the leading eigenvalue will be given by:

$$\lambda_{max} = \frac{1}{2}\left(A + \sqrt{A^2 + B + C}\right) \tag{S80}$$

Where:

$$A = \zeta\mu\sigma_{f_1} + (1-\zeta)\nu\sigma_{m_1} \tag{S81a}$$

$$B = 4\zeta(1-\mu)\sigma_{\mathrm{f}_2}\left(\alpha\tau_{\mathrm{f}} + (1-\alpha)\tau_{\mathrm{m}}\right) \quad \text{(S81b)}$$

$$C = 4(1-\zeta)(1-\nu)\sigma_{\mathrm{m}_2}\left((1-\beta)\tau_{\mathrm{f}} + \beta\tau_{\mathrm{m}}\right) \quad \text{(S81c)}$$

And:

$$\sigma_{\mathrm{f}_1} = \frac{\omega_{\{\mathrm{f}_1,1\}}}{\omega_{\{\mathrm{f}_1,0\}}}, \frac{\omega_{\{\mathrm{f}_1,0\}}}{\omega_{\{\mathrm{f}_1,1\}}} \quad \text{(S82a)}$$

$$\sigma_{\mathrm{f}_2} = \frac{\omega_{\{\mathrm{f}_2,1\}}}{\omega_{\{\mathrm{f}_2,0\}}}, \frac{\omega_{\{\mathrm{f}_2,0\}}}{\omega_{\{\mathrm{f}_2,1\}}} \quad \text{(S82b)}$$

$$\sigma_{\mathrm{m}_1} = \frac{\omega_{\{\mathrm{m}_1,1\}}}{\omega_{\{\mathrm{m}_1,0\}}}, \frac{\omega_{\{\mathrm{m}_1,0\}}}{\omega_{\{\mathrm{m}_1,1\}}} \quad \text{(S82c)}$$

$$\sigma_{\mathrm{m}_2} = \frac{\omega_{\{\mathrm{m}_2,1\}}}{\omega_{\{\mathrm{m}_2,0\}}}, \frac{\omega_{\{\mathrm{m}_2,0\}}}{\omega_{\{\mathrm{m}_2,1\}}} \quad \text{(S82d)}$$

$$\tau_{\mathrm{f}} = \frac{\Omega_{\{\mathrm{f},1\}}}{\Omega_{\{\mathrm{f},0\}}}, \frac{\Omega_{\{\mathrm{f},0\}}}{\Omega_{\{\mathrm{f},1\}}} \quad \text{(S82e)}$$

$$\tau_{\mathrm{m}} = \frac{\Omega_{\{\mathrm{m},1\}}}{\Omega_{\{\mathrm{m},0\}}}, \frac{\Omega_{\{\mathrm{m},0\}}}{\Omega_{\{\mathrm{m},1\}}} \quad \text{(S82f)}$$

##### 3.3.2 Diploid

For the haploid system, the leading eigenvalue will be given by:

$$\lambda_{max} = \frac{1}{4}\left(A + \sqrt{A^2 + B + C}\right) \quad \text{(S83)}$$

Where:

$$A = \mu\sigma_{\mathrm{f}_1} + \nu\sigma_{\mathrm{m}_1} \quad \text{(S84a)}$$

$$B = 8(1-\mu)\sigma_{\mathrm{f}_2}\left(\alpha\tau_{\mathrm{f}} + (1-\alpha)\tau_{\mathrm{m}}\right) \quad \text{(S84b)}$$

$$C = 8(1-\nu)\sigma_{\mathrm{m}_2}\left((1-\beta)\tau_{\mathrm{f}} + \beta\tau_{\mathrm{m}}\right) \quad \text{(S84c)}$$

And:

$$\sigma_{\mathrm{f}_1} = \left\{\frac{\omega_{\{\mathrm{f}_1,01\}}}{\omega_{\{\mathrm{f}_1,00\}}}, \frac{\omega_{\{\mathrm{f}_1,10\}}}{\omega_{\{\mathrm{f}_1,00\}}}\right\}, \left\{\frac{\omega_{\{\mathrm{f}_1,01\}}}{\omega_{\{\mathrm{f}_1,11\}}}, \frac{\omega_{\{\mathrm{f}_1,10\}}}{\omega_{\{\mathrm{f}_1,11\}}}\right\} \quad \text{(S85a)}$$

$$\sigma_{\mathrm{f}_2} = \left\{\frac{\omega_{\{\mathrm{f}_2,01\}}}{\omega_{\{\mathrm{f}_2,00\}}}, \frac{\omega_{\{\mathrm{f}_2,10\}}}{\omega_{\{\mathrm{f}_2,00\}}}\right\}, \left\{\frac{\omega_{\{\mathrm{f}_2,01\}}}{\omega_{\{\mathrm{f}_2,11\}}}, \frac{\omega_{\{\mathrm{f}_2,10\}}}{\omega_{\{\mathrm{f}_2,11\}}}\right\} \quad \text{(S85b)}$$

$$\sigma_{\mathrm{m}_1} = \left\{\frac{\omega_{\{\mathrm{m}_1,01\}}}{\omega_{\{\mathrm{m}_1,00\}}}, \frac{\omega_{\{\mathrm{m}_1,10\}}}{\omega_{\{\mathrm{m}_1,00\}}}\right\}, \left\{\frac{\omega_{\{\mathrm{m}_1,01\}}}{\omega_{\{\mathrm{m}_1,11\}}}, \frac{\omega_{\{\mathrm{m}_1,10\}}}{\omega_{\{\mathrm{m}_1,11\}}}\right\} \quad \text{(S85c)}$$

$$\sigma_{\mathrm{m}_2} = \left\{\frac{\omega_{\{\mathrm{m}_2,01\}}}{\omega_{\{\mathrm{m}_2,00\}}}, \frac{\omega_{\{\mathrm{m}_2,10\}}}{\omega_{\{\mathrm{m}_2,00\}}}\right\}, \left\{\frac{\omega_{\{\mathrm{m}_2,01\}}}{\omega_{\{\mathrm{m}_2,11\}}}, \frac{\omega_{\{\mathrm{m}_2,10\}}}{\omega_{\{\mathrm{m}_2,11\}}}\right\} \quad \text{(S85d)}$$

$$\tau_{\mathrm{f}} = \left\{\frac{\Omega_{\{\mathrm{f},01\}}}{\Omega_{\{\mathrm{f},00\}}}, \frac{\Omega_{\{\mathrm{f},10\}}}{\Omega_{\{\mathrm{f},00\}}}\right\}, \left\{\frac{\Omega_{\{\mathrm{f},01\}}}{\Omega_{\{\mathrm{f},11\}}}, \frac{\Omega_{\{\mathrm{f},10\}}}{\Omega_{\{\mathrm{f},11\}}}\right\} \quad \text{(S85e)}$$

$$\tau_{\mathrm{m}} = \left\{\frac{\Omega_{\{\mathrm{m},01\}}}{\Omega_{\{\mathrm{m},00\}}}, \frac{\Omega_{\{\mathrm{m},10\}}}{\Omega_{\{\mathrm{m},00\}}}\right\}, \left\{\frac{\Omega_{\{\mathrm{m},01\}}}{\Omega_{\{\mathrm{m},11\}}}, \frac{\Omega_{\{\mathrm{m},10\}}}{\Omega_{\{\mathrm{m},11\}}}\right\} \quad \text{(S85f)}$$

#### 3.4 Invasion conditions

We can calculate the invasion conditions for various types of sexually antagonistic allele by first substituting in the values from the fitness scheme in Tables S1 and S2), and then solving for when $\lambda_{\max} > 1$.

##### 3.4.1 Sexual antagonism over fecundity

For a sexually antagonistic allele that affects fecundity in both sexes ($s_\mathrm{f}^\mathrm{s} = s_\mathrm{m}^\mathrm{s} = 0, s_\mathrm{f}^\mathrm{r} = s_\mathrm{f}, s_\mathrm{m}^\mathrm{r} = s_\mathrm{m}$), then a female beneficial will invade under the haploid system when:

$$s_\mathrm{m} < \frac{\zeta s_\mathrm{f}}{(1-\zeta)(1-s_\mathrm{f})} \tag{S86}$$

And fixes when:

$$s_\mathrm{m} < \frac{\zeta s_\mathrm{f}}{(1-\zeta)+\zeta s_\mathrm{f}} \tag{S87}$$

And under the diploid system will invade when:

$$s_\mathrm{m} < \frac{(1-h_\mathrm{f})s_\mathrm{f}}{h_\mathrm{m}(1-s_\mathrm{f})} \tag{S88}$$

and fixes when:

$$s_\mathrm{m} < \frac{h_\mathrm{f}s_\mathrm{f}}{h_\mathrm{f}s_\mathrm{f}+(1-h_\mathrm{m})} \tag{S89}$$

Which under weak selection we can rewrite for the haploid system as:

$$c_\mathrm{m}^\mathrm{r}(s_\mathrm{m}) < c_\mathrm{f}^\mathrm{r}(s_\mathrm{f}) \tag{S90}$$

And:

$$c_\mathrm{m}^\mathrm{r}(s_\mathrm{m}) < c_\mathrm{f}^\mathrm{r}(s_\mathrm{f}) \tag{S91}$$

And for the diploid system as:

$$c_\mathrm{m}^\mathrm{r}(h_\mathrm{m}s_\mathrm{m}) < c_\mathrm{f}^\mathrm{r}((1-h_\mathrm{f})s_\mathrm{f}) \tag{S92}$$

And:

$$c_\mathrm{m}^\mathrm{r}((1-h_\mathrm{m})s_\mathrm{m}) < c_\mathrm{f}^\mathrm{r}(h_\mathrm{f}s_\mathrm{f}) \tag{S93}$$

With the reverse of these for a male beneficial allele.

##### 3.4.2 Sexual antagonism over survival

For a sexually antagonistic allele that affects survival in both sexes ($s_\mathrm{f}^\mathrm{r} = s_\mathrm{m}^\mathrm{r} = 0, s_\mathrm{f}^\mathrm{s} = s_\mathrm{f}, s_\mathrm{m}^\mathrm{s} = s_\mathrm{m}$), then the female beneficial allele invades under haploidy when:

$$s_\mathrm{m} < \frac{\big(\alpha\zeta(1-\mu)+(1-\beta)(1-\zeta)(1-\nu)\big)\,s_\mathrm{f}}{\big((1-\alpha)\zeta(1-\mu)+\beta(1-\zeta)(1-\nu)\big)\,(1-s_\mathrm{f})} \tag{S94}$$

And fixes when:

$$s_\mathrm{m} < \frac{\big(\alpha\zeta(1-\mu)+(1-\beta)(1-\zeta)(1-\nu)\big)\,s_\mathrm{f}}{(1-\alpha)\zeta(1-\mu)+\beta(1-\zeta)(1-\nu)+\big(\alpha\zeta(1-\mu)+(1-\beta)(1-\zeta)(1-\nu)\big)\,s_\mathrm{f}} \tag{S95}$$

And for diploidy, the invasion condition will be:

$$s_\mathrm{m} < \frac{(1-h_\mathrm{f})s_\mathrm{f}(\alpha(1-\mu)+(1-\beta)(1-\nu))}{h_\mathrm{m}(1-s_\mathrm{f})((1-\alpha)(1-\mu)+\beta(1-\nu))} \tag{S96}$$

and the fixation condition will be:

$$s_\mathrm{m} < \frac{h_\mathrm{f}s_\mathrm{f}\big(\alpha(1-\mu)+(1-\beta)(1-\nu)\big)}{h_\mathrm{f}s_\mathrm{f}\big(\alpha(1-\mu)+(1-\beta)(1-\nu)\big)+(1-h_\mathrm{m})\big((1-\alpha)(1-\mu)+\beta(1-\nu)\big)} \tag{S97}$$

Which again under weak selection can be rewritten. For haploidy, invasion and fixation are given by:

$$c_{\mathrm{m}}^{\mathrm{s}}\,(s_{\mathrm{m}}) < c_{\mathrm{f}}^{\mathrm{s}}\,(s_{\mathrm{f}}) \tag{S98}$$

Where $c_{\mathrm{f}}^{\mathrm{s}}$ and $c_{\mathrm{m}}^{\mathrm{s}}$ are the class reproductive values of males and females through survival. And for diploidy, invasion will be:

$$c_{\mathrm{m}}^{\mathrm{s}}\,(h_{\mathrm{m}} s_{\mathrm{m}}) < c_{\mathrm{f}}^{\mathrm{s}}\,((1-h_{\mathrm{f}})\,s_{\mathrm{f}}) \tag{S99}$$

With the fixation condition as:

$$c_{\mathrm{m}}^{\mathrm{s}}\,((1-h_{\mathrm{m}})\,s_{\mathrm{m}}) < c_{\mathrm{f}}^{\mathrm{s}}\,(h_{\mathrm{f}} s_{\mathrm{f}}) \tag{S100}$$

The conditions for a male beneficial allele will be the reverse of these.

##### 3.4.3 Sexual antagonism over survival and fecundity

Now, we can consider sexual antagonism that manifests in different functions (i.e. survival and fecundity). The condition for a sexually antagonistic allele which confers a benefit to female fecundity, but cost to male survival (i.e. $s_{\mathrm{f}}^{\mathrm{s}} = s_{\mathrm{m}}^{\mathrm{r}} = 0, s_{\mathrm{f}}^{\mathrm{r}} = s_{\mathrm{f}}, s_{\mathrm{m}}^{\mathrm{s}} = s_{\mathrm{m}}$) will invade the population under haploidy is:

$$s_{\mathrm{m}} < \frac{\zeta s_{\mathrm{f}}}{(1-\alpha)\zeta(1-\mu) + \beta(1-\zeta)(1-\nu)\,(1-s_{\mathrm{f}})} \tag{S101}$$

and will fix when:

$$s_{\mathrm{m}} < \frac{\zeta s_{\mathrm{f}}}{\beta(1-\zeta)(1-\nu) + (1-\alpha)(\zeta(1-\mu))\,(1-s_{\mathrm{f}})} \tag{S102}$$

And will invade under diploidy when:

$$s_{\mathrm{m}} < \frac{(1-h_{\mathrm{f}})\,s_{\mathrm{f}}}{h_{\mathrm{m}}\left((1-\alpha)(1-\mu)(1-h_{\mathrm{f}} s_{\mathrm{f}}) + \beta(1-\nu)(1-s_{\mathrm{f}})\right)} \tag{S103}$$

And will reach fixation when:

$$s_{\mathrm{m}} < \frac{h_{\mathrm{f}} s_{\mathrm{f}}}{h_{\mathrm{f}} s_{\mathrm{f}} + (1-h_{\mathrm{m}})\left((1-\beta)(1-\nu) + \alpha(1-\mu)(1-h_{\mathrm{f}} s_{\mathrm{f}})\right)} \tag{S104}$$

Again, under weak selection we can simplify and rewrite these conditions. For haploidy, invasion and fixation are:

$$c_{\mathrm{m}}^{\mathrm{s}}\,(s_{\mathrm{m}}) < c_{\mathrm{f}}^{\mathrm{r}}\,(s_{\mathrm{f}}) \tag{S105}$$

And for diploidy, invasion will be:

$$c_{\mathrm{m}}^{\mathrm{s}}\,(h_{\mathrm{m}} s_{\mathrm{m}}) < c_{\mathrm{f}}^{\mathrm{r}}\,((1-h_{\mathrm{f}})\,s_{\mathrm{f}}) \tag{S106}$$

And fixation will be:

$$c_{\mathrm{m}}^{\mathrm{s}}\,((1-h_{\mathrm{m}})\,s_{\mathrm{m}}) < c_{\mathrm{f}}^{\mathrm{r}}\,(h_{\mathrm{f}} s_{\mathrm{f}}) \tag{S107}$$

Conversely, if the allele is beneficial to female survival, but costly to male fecundity (i.e. $s_{\mathrm{f}}^{\mathrm{r}} = s_{\mathrm{m}}^{\mathrm{s}} = 0, s_{\mathrm{f}}^{\mathrm{s}} = s_{\mathrm{f}}, s_{\mathrm{m}}^{\mathrm{r}} = s_{\mathrm{m}}$) then the condition for the female beneficial allele to invade will, under haploidy, be:

$$s_{\mathrm{m}} < \frac{s_{\mathrm{f}}(\alpha\zeta(1-\mu) + (1-\beta)(1-\zeta)(1-\nu))}{(1-\zeta) - (1-\zeta)(\beta(1-\nu)+\nu)\,s_{\mathrm{f}}} \tag{S108}$$

And to fix will be:

$$s_{\mathrm{m}} < \frac{\left(\alpha\zeta(1-\mu) + (1-\beta)(1-\zeta)(1-\nu)\right)s_{\mathrm{f}}}{(1-\zeta) + \alpha\zeta(1-\mu)\,s_{\mathrm{f}}} \tag{S109}$$

And for diploids, the condition for invasion will be:

$$s_{\mathrm{m}} < \frac{(1-h_{\mathrm{f}})s_{\mathrm{f}}\big(\alpha(1-\mu)+(1-\beta)(1-\nu)\big)}{h_{\mathrm{m}}(1-s_{\mathrm{f}}+(1-h_{\mathrm{f}})s_{\mathrm{f}}(1-\beta)(1-\nu))} \tag{S110}$$

With the fixation condition:

$$s_{\mathrm{m}} < \frac{h_{\mathrm{f}}s_{\mathrm{f}}(\alpha(1-\mu)+(1-\beta)(1-\nu))}{(1-h_{\mathrm{m}})+s_{\mathrm{f}}(\alpha(1-\mu)h_{\mathrm{f}}+(1-\beta)(1-\nu)h_{\mathrm{f}}h_{\mathrm{m}})} \tag{S111}$$

Which under weak selection these can be rewritten. For haploidy invasion and fixation are:

$$c_{\mathrm{m}}^{\mathrm{r}}(s_{\mathrm{m}}) < c_{\mathrm{f}}^{\mathrm{s}}(s_{\mathrm{f}}) \tag{S112}$$

And for diploidy, the invasion condition is given by:

$$c_{\mathrm{m}}^{\mathrm{r}}(h_{\mathrm{m}}s_{\mathrm{m}}) < c_{\mathrm{f}}^{\mathrm{s}}((1-h_{\mathrm{f}})s_{\mathrm{f}}) \tag{S113}$$

And with the associated fixation condition of:

$$c_{\mathrm{m}}^{\mathrm{r}}((1-h_{\mathrm{m}})s_{\mathrm{m}}) < c_{\mathrm{f}}^{\mathrm{s}}(h_{\mathrm{f}}s_{\mathrm{f}}) \tag{S114}$$

Once again, we can recover the conditions for a male beneficial allele to invade and fix, by reversing the signs of these equations.

##### 3.5 Within sex trade-offs over survival and fecundity

We can also consider some non-sexually antagonistic trade-offs. Here we instead use the fitness schemes described in Tables S3 and S4, where we instead consider an allele which confers a fecundity benefit, but survival cost. First, we consider the case where it has the same fitness effects in both sexes (i.e. $s_{\mathrm{f}}^{\mathrm{r}} = s_{\mathrm{m}}^{\mathrm{r}} = s_{\mathrm{r}}, s_{\mathrm{f}}^{\mathrm{s}} = s_{\mathrm{m}}^{\mathrm{s}} = s_{\mathrm{s}}$).In this case, the condition for invasion in the haploid system will be:

$$s_{\mathrm{s}} < \frac{s_{\mathrm{r}}}{\zeta(1-\mu)+(1-\zeta)(1-\nu)} \tag{S115}$$

and for fixation:

$$s_{\mathrm{s}} < \frac{s_{\mathrm{r}}}{\zeta(1-\mu)+(1-\zeta)(1-\nu)+(\zeta\mu+(1-\zeta)\nu)s_{\mathrm{r}}} \tag{S116}$$

And in the diploid system, the condition for invasion is:

$$s_{\mathrm{s}} < \frac{2(1-h_{\mathrm{r}})s_{\mathrm{r}}}{(2-\mu-\nu)h_{\mathrm{s}}-(2-\mu-\nu)h_{\mathrm{r}}h_{\mathrm{s}}s_{\mathrm{r}}} \tag{S117}$$

And for fixation:

$$s_{\mathrm{s}} < \frac{2h_{\mathrm{r}}s_{\mathrm{r}}}{(2-\mu-\nu)(1-h_{\mathrm{s}})+\big((\mu+\nu)h_{\mathrm{r}}+(2-\mu-\nu)h_{\mathrm{r}}h_{\mathrm{s}}\big)s_{\mathrm{r}}} \tag{S118}$$

Assuming weak selection, we can simplify and rewrite. For haploidy, the invasion and fixation conditions become:

$$\left(c_{\mathrm{f}}^{\mathrm{s}}+c_{\mathrm{m}}^{\mathrm{s}}\right)(s_{\mathrm{s}}) < \left(c_{\mathrm{f}}^{\mathrm{r}}+c_{\mathrm{m}}^{\mathrm{r}}\right)(s_{\mathrm{r}}) \tag{S119}$$

And for diploidy, the invasion condition becomes:

$$\left(c_{\mathrm{f}}^{\mathrm{s}}+c_{\mathrm{m}}^{\mathrm{s}}\right)(h_{\mathrm{s}}s_{\mathrm{s}}) < \left(c_{\mathrm{f}}^{\mathrm{r}}+c_{\mathrm{m}}^{\mathrm{r}}\right)((1-h_{\mathrm{f}})s_{\mathrm{r}}) \tag{S120}$$

With the fixation condition of:

$$\left(c_{\mathrm{f}}^{\mathrm{s}}+c_{\mathrm{m}}^{\mathrm{s}}\right)\left((1-h_{\mathrm{s}})s_{\mathrm{s}}\right)<\left(c_{\mathrm{f}}^{\mathrm{r}}+c_{\mathrm{m}}^{\mathrm{r}}\right)\left(h_{\mathrm{f}}s_{\mathrm{r}}\right) \tag{S121}$$

We can also consider such trade-offs being exclusively within one sex. In females, under the haploid system, the allele will invade provided:

$$s_{\mathrm{s}}<\frac{\zeta s_{\mathrm{r}}}{\alpha\zeta(1-\mu)+(1-\beta)(1-\zeta)(1-\nu)(1-s_{\mathrm{r}})} \tag{S122}$$

And will fix provided:

$$s_{\mathrm{s}}<\frac{\zeta s_{\mathrm{r}}}{\alpha(\zeta(1-\mu))+(1-\beta)(1-\zeta)(1-\nu)+\zeta(1-\alpha(1-\mu))s_{\mathrm{r}}} \tag{S123}$$

And for the diploid system, the allele will invade when:

$$s_{\mathrm{s}}<\frac{(1-h_{\mathrm{r}})s_{\mathrm{r}}}{\left(\alpha(1-\mu)+(1-\beta)(1-\nu)\right)h_{\mathrm{s}}-\left((1-\beta)(1-\nu)+\alpha(1-\mu)h_{\mathrm{r}}\right)h_{\mathrm{s}}s_{\mathrm{r}}} \tag{S124}$$

And will fix when:

$$s_{\mathrm{s}}<\frac{h_{\mathrm{r}}s_{\mathrm{r}}}{\left(\alpha(1-\mu)+(1-\beta)(1-\nu)\right)(1-h_{\mathrm{s}})+\left(1-\alpha(1-\mu)(1-h_{\mathrm{s}})\right)h_{\mathrm{r}}s_{\mathrm{r}}} \tag{S125}$$

Under weak selection, the haploid condition for invasion and fixation may be written as:

$$c_{\mathrm{f}}^{\mathrm{s}}\left(s_{\mathrm{s}}\right)<c_{\mathrm{f}}^{\mathrm{r}}\left(s_{\mathrm{r}}\right) \tag{S126}$$

And for diploidy, the invasion condition becomes:

$$c_{\mathrm{f}}^{\mathrm{s}}\left(h_{\mathrm{s}}s_{\mathrm{s}}\right)<c_{\mathrm{f}}^{\mathrm{r}}\left((1-h_{\mathrm{f}})s_{\mathrm{r}}\right) \tag{S127}$$

With the fixation condition of:

$$c_{\mathrm{f}}^{\mathrm{s}}\left((1-h_{\mathrm{s}})s_{\mathrm{s}}\right)<c_{\mathrm{f}}^{\mathrm{r}}\left(h_{\mathrm{f}}s_{\mathrm{r}}\right) \tag{S128}$$

Similarly, looking at males, an allele which provides a fecundity benefit but survival cost will invade under haploidy provided:

$$s_{\mathrm{s}}<\frac{(1-\zeta)s_{\mathrm{r}}}{\beta(1-\zeta)(1-\nu)+(1-\alpha)\zeta(1-\mu)(1-s_{\mathrm{r}})} \tag{S129}$$

And will fix provided:

$$\frac{(1-\zeta)s_{\mathrm{r}}}{(1-\alpha)\zeta(1-\mu)+\beta(1-\zeta)(1-\nu)+(1-\zeta)(1-\beta(1-\nu))s_{\mathrm{r}}} \tag{S130}$$

And under diploidy, invasion will occur when:

$$s_{\mathrm{s}}<\frac{(1-h_{\mathrm{r}})s_{\mathrm{r}}}{\left((1-\alpha)(1-\mu)+\beta(1-\nu)\right)h_{\mathrm{s}}-\left((1-\alpha)(1-\mu)+(\beta(1-\nu))h_{\mathrm{r}}\right)h_{\mathrm{s}}s_{\mathrm{r}}} \tag{S131}$$

And will fix when:

$$s_{\mathrm{s}}<\frac{h_{\mathrm{r}}s_{\mathrm{r}}}{(1-h_{\mathrm{s}})\left((1-\alpha)(1-\mu)+\beta(1-\nu)\right)+\left(1-(\beta(1-\nu))(1-h_{\mathrm{s}})\right)h_{\mathrm{r}}s_{\mathrm{r}}} \tag{S132}$$

Under weak selection these conditions may be rewritten. For haploidy, invasion and fixation are given by:

$$c_{\mathrm{m}}^{\mathrm{s}}\left(s_{\mathrm{s}}\right)<c_{\mathrm{m}}^{\mathrm{r}}\left(s_{\mathrm{r}}\right) \tag{S133}$$

And for diploidy, invasion will be given by:

$$c_{\mathrm{m}}^{\mathrm{s}}\left(h_{\mathrm{s}}s_{\mathrm{s}}\right)<c_{\mathrm{m}}^{\mathrm{r}}\left((1-h_{\mathrm{f}})s_{\mathrm{r}}\right) \tag{S134}$$

And fixation by:

$$c_{\mathrm{m}}^{\mathrm{s}}\left((1-h_{\mathrm{s}})s_{\mathrm{s}}\right)<c_{\mathrm{m}}^{\mathrm{r}}\left(h_{\mathrm{f}}s_{\mathrm{r}}\right) \tag{S135}$$

#### 3.6 Tables and Figures

Table S1: **Fitness scheme for haploid model of sexual antagonism**.

| | $\omega_0$ | $\omega_1$ | $\Omega_0$ | $\Omega_1$ |
|---|---|---|---|---|
| $F_1$ | $1-s_f^r$ | 1 | $1-s_f^s$ | 1 |
| $F_2$ | $1-s_f^r$ | 1 | n/a | n/a |
| $M_1$ | 1 | $1-s_m^r$ | 1 | $1-s_m^s$ |
| $M_2$ | 1 | $1-s_m^r$ | n/a | n/a |

Table S2: **Fitness scheme for diploid model of sexual antagonism**.

| | $\omega_{00}$ | $\omega_{01}/\omega_{10}$ | $\omega_{11}$ | $\Omega_{00}$ | $\Omega_{01}/\Omega_{10}$ | $\Omega_{11}$ |
|---|---|---|---|---|---|---|
| $F_1$ | $1-s_f^r$ | $1-h_f s_f^r$ | 1 | $1-s_f^s$ | $1-h_f s_f^s$ | 1 |
| $F_2$ | $1-s_f^r$ | $1-h_f s_f^r$ | 1 | n/a | n/a | n/a |
| $M_1$ | 1 | $1-h_m s_m^r$ | $1-s_m^r$ | 1 | $1-h_m s_m^s$ | $1-s_m^s$ |
| $M_2$ | 1 | $1-h_m s_m^r$ | $1-s_m^r$ | n/a | n/a | n/a |

Table S3: **Fitness scheme for haploid model of fecundity/survival trade-offs**.

| | $\omega_0$ | $\omega_1$ | $\Omega_0$ | $\Omega_1$ |
|---|---|---|---|---|
| $F_1$ | $1-s_f^r$ | 1 | 1 | $1-s_f^s$ |
| $F_2$ | $1-s_f^r$ | 1 | n/a | n/a |
| $M_1$ | $1-s_m^r$ | 1 | 1 | $1-s_m^s$ |
| $M_2$ | $1-s_m^r$ | 1 | n/a | n/a |

Table S4: **Fitness scheme for diploid model of fecundity/survival trade-offs**.

| | $\omega_{00}$ | $\omega_{01}/\omega_{10}$ | $\omega_{11}$ | $\Omega_{00}$ | $\Omega_{01}/\Omega_{10}$ | $\Omega_{11}$ |
|---|---|---|---|---|---|---|
| $F_1$ | $1-s_{\mathrm{f}}^{\mathrm{r}}$ | $1-h_{\mathrm{r}}s_{\mathrm{f}}^{\mathrm{r}}$ | 1 | 1 | $1-h_{\mathrm{d}}s_{\mathrm{f}}^{\mathrm{s}}$ | $1-s_{\mathrm{f}}^{\mathrm{s}}$ |
| $F_2$ | $1-s_{\mathrm{f}}^{\mathrm{r}}$ | $1-h_{\mathrm{r}}s_{\mathrm{f}}^{\mathrm{r}}$ | 1 | n/a | n/a | n/a |
| $M_1$ | $1-s_{\mathrm{m}}^{\mathrm{r}}$ | $1-h_{\mathrm{r}}s_{\mathrm{m}}^{\mathrm{r}}$ | 1 | 1 | $1-h_{\mathrm{s}}s_{\mathrm{m}}^{\mathrm{s}}$ | $1-s_{\mathrm{m}}^{\mathrm{s}}$ |
| $M_2$ | $1-s_{\mathrm{m}}^{\mathrm{r}}$ | $1-h_{\mathrm{r}}s_{\mathrm{m}}^{\mathrm{r}}$ | 1 | n/a | n/a | n/a |

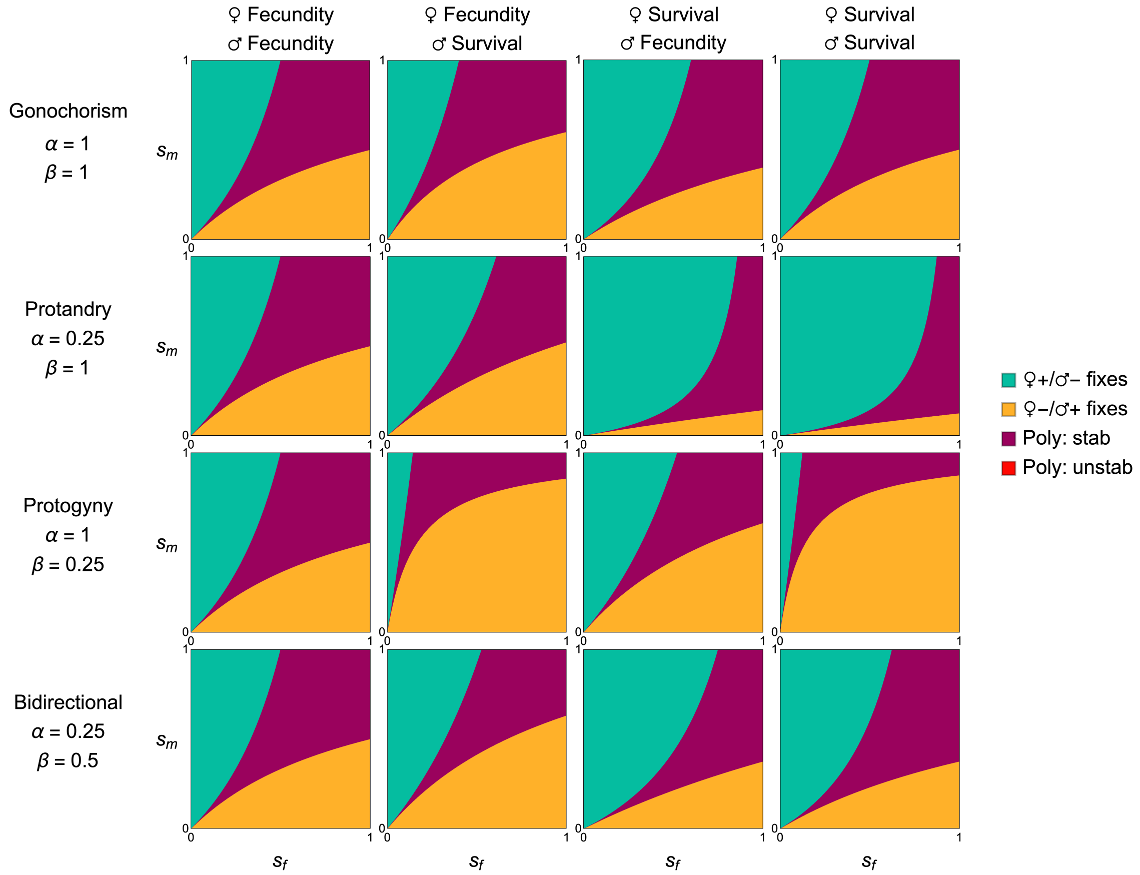


Figure S2: **Invasion conditions for an allele affecting various types of sexually antagonistic trade-off under reversals of dominance.** Trade-offs plotted include: fecundity in both sexes, fecundity in females and survival in males, survival in females and fecundity in males, and survival in both sexes. And for various types of sexual system including gonochorism, protandry, protogyny and bidirectional sex change. The meaning of key parameters are summarised in Figure S1. For all plots we assume that $\mu = \nu = 1/3$, and $h_{\mathrm{f}} = h_{\mathrm{m}} = 1/4$.

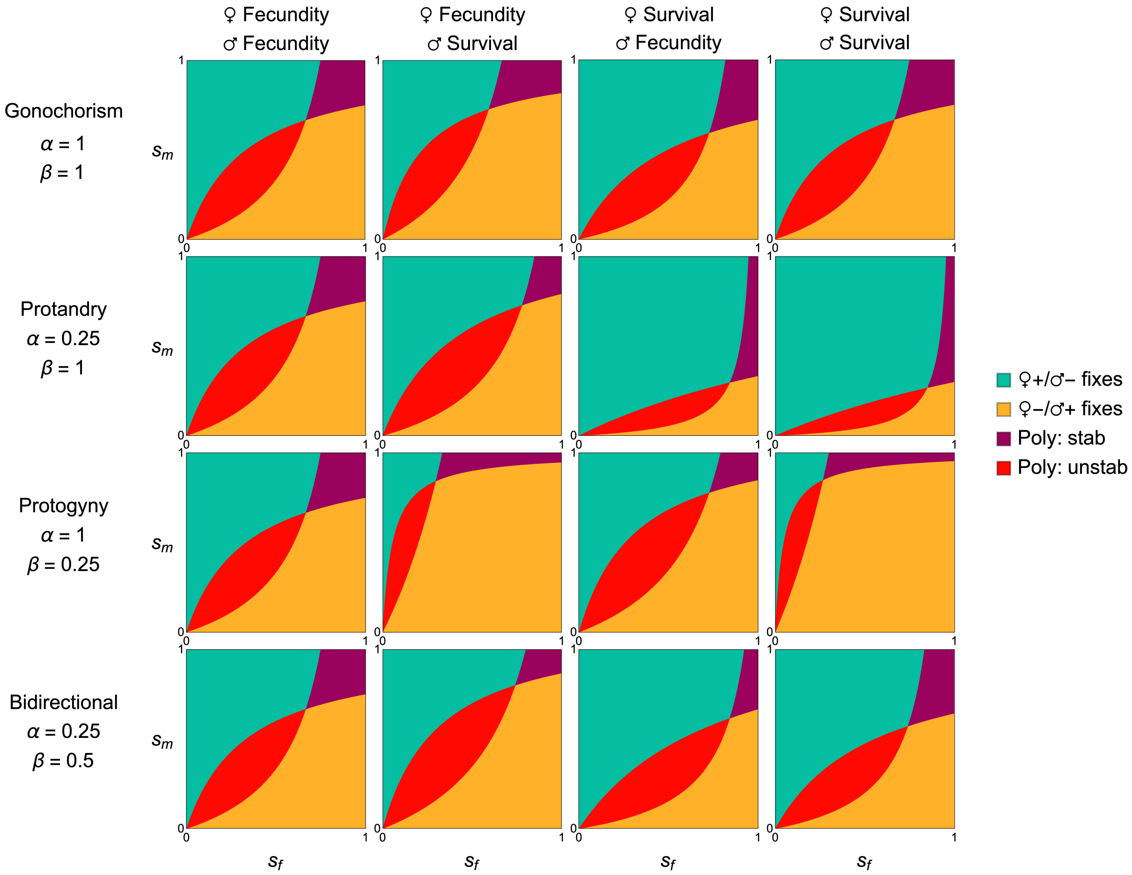


Figure S3: **Invasion conditions for an allele affecting various types of sexually antagonistic trade-off under reversals of dominance.** Trade-offs plotted include: fecundity in both sexes, fecundity in females and survival in males, survival in females and fecundity in males, and survival in both sexes. And for various types of sexual system including gonochorism, protandry, protogyny and bidirectional sex change. The meaning of key parameters are summarised in Figure S1. For all plots we assume that $\mu = \nu = 1/3$, and $h_\mathrm{f} = h_\mathrm{m} = 3/4$.
